# The Aging Synapse: An Integrated Proteomic And Transcriptomic Analysis

**DOI:** 10.1101/2024.12.18.629115

**Authors:** Cinzia Caterino, Martino Ugolini, William Durso, Kristina Jevdokimenko, Marco Groth, Konstantin Riege, Matthias Görlach, Eugenio Fornasiero, Alessandro Ori, Steve Hoffmann, Alessandro Cellerino

## Abstract

An important hallmark of aging is the loss of proteostasis, which can lead to the formation of protein aggregates and mitochondrial dysfunction in neurons. Although it is well known that protein synthesis is finely regulated in the brain, especially at synapses, where mRNAs are locally translated in activity-dependent manner, little is known as to the changes in the synaptic proteome and transcriptome during aging. Therefore, this work aims to elucidate the relationship between transcriptome and proteome at soma and synaptic level during aging.

Proteomic and transcriptomic data analysis reveal that, in young animals, proteins and transcripts are correlated and synaptic regulation is driven by changes in the soma. During aging, there is a decoupling between transcripts and proteins and between somatic and synaptic compartments. Furthermore, soma-synapse gradient of ribosomal genes changes upon aging, i.e. ribosomal transcripts are less abundant and ribosomal proteins are more abundant in synaptic compartment of old mice with respect to younglings. Additionally, transcriptomics data highlight a difference in the splicing of certain synaptic mRNA with aging. Taken together, our data provide a valuable resource for the study of the aging synapse.

## INTRODUCTION

Brain aging is characterized by progressive reduction of cortical grey matter volume (Fjell et al., 2009; Lemaître et al., 2005, 2012; Raz et al., 2005). Surprisingly, cortical shrinkage is not correlated with significant neuronal death during normal brain aging but represents a pathological feature associated with neurodegenerative diseases (Morrison & Hof, 2007; Pakkenberg et al., 2003). Cortical shrinkage must then be mainly the result of changes in the network of neuronal connections (neuropil) and indeed aged pyramidal cortical neurons are affected by reduction of synaptic spines (Dickstein et al., 2007, 2013) and imbalances in neurotransmitter signaling (Leventhal et al., 2003; Wang et al., 2011). These synaptic changes are likely the cause of age-dependent deficits in functions such as perceptual speed and working memory (Buckner, 2004; Nyberg et al., 2012).

Neurons are cells of a peculiar morphology that comprises complex and extensive neurites. Neurites can have a length that is orders of magnitude larger than the soma diameter and most of neuronal cytoplasm is contained in neurites. Dispatching somatically-synthesized proteins to distant compartments is a demanding process and it has become increasingly clear that many transcripts coding for synaptic proteins are transported to distant sites (von Kügelgen & Chekulaeva, 2020 and references therein) and that most of neuritic proteome is synthesized locally (Zappulo et al., 2017). Local synaptic translation is indispensable both for synapse maintenance and for activity-dependent synaptic plasticity both during development and in adulthood (Cioni et al., 2018; Glock et al., 2017; Holt & Schuman, 2013; Holt et al., 2019; Sun et al., 2021). Transcripts coding for synaptic proteins are subject to alternative splicing and are enriched in specific motifs that support local translation. Neuronal aging is known to affect splicing at a global level and altered splicing is a likely driving factor of Alzheimer’s disease (Raj et al., 2018).

Although synaptic translation has been extensively analyzed in relation to several physiological and pathological mechanisms (Piol et al., 2023; Seo et al., 2022; Cagnetta et al., 2023), no one has so far provided a comprehensive view of the changes that occur at the synaptic level with age. Albeit there is no lack of studies on the modification of the synaptic transcriptome during ageing (Chen et al., 2017a; Dillman et al., 2017), here we provide the first insight of how transcriptome and proteome change together in the aging synapse. Furthermore, in this work we analyze the effects of aging on differential splicing in the local synaptic proteome. To this end, we analyzed the RNA extracted from synaptosomes obtained from cerebral cortices of C57BL/6J mice of 3 weeks, 5 months and 18 months of age. These three time points span three pivotal stages of brain development: at 3 weeks corticogenesis is completed but synaptogenesis is ongoing (M. Li et al., 2010) and cortical plasticity is likely at its peak (Gordon & Stryker, 1996). At 5 months the mouse is considered mature adult, and this represents an intermediate time point between development and aging. Mice at 18 months of age already suffer from cognitive decay (Fukushima et al., 2008; Peleg et al., 2010) but age-dependent mortality for the BL6 strain is still marginal, with survivorship of around 80% (Flurkey et al., 2007). Synaptosomes are artificial, membranous sacs that contain synaptic components and are generated by subcellular fractionation of homogenized or ground-up nerve tissue, because the lipid bilayers naturally reseal together after the axon terminals are torn off by the physical shearing force of homogenization. Synaptosomes contain the complete presynaptic terminal, including mitochondria and synaptic vesicles, along with the postsynaptic membrane and the postsynaptic density (PSD)(Daniel et al., 2012). This preparation has been widely used for biochemical characterization of synapses and to study mechanisms of neurotransmitter release (Trebesova & Grilli, 2023;Garcia-Sanz et al., 2001). Here we employ RNA-seq, proteomics and sequencing of ribosome-associated RNAs to obtain the first multi-omics analysis of the aging synapse.

## RESULTS AND DISCUSSION

### Synaptosomes enrichment from mouse cortex during aging

We extracted cortex-derived synaptosome (SYN) samples from young (3 weeks old), adult (5 months old) and old (18 months old) C57BL/6J mice (Table S1). From each sample, both RNA and proteins were purified. The enrichment of synaptic components in SYN samples was verified through Western Blot and quantitative Real-Time PCR (qPCR) (**Figure S1A-G**).

We then performed depletion of rRNAs and single-end RNA-Seq on SYN and total homogenate (TH) samples, obtaining an average number of reads per sample of 25 ± 1 and 20 ± 3 millions, respectively. The distribution of reads mapping to different genomic features differs between SYN and TH, with higher percentage of reads mapping to intronic sequences in TH samples as compared to SYN samples (**Figure S1H**), as expected due to the presence of pre-mRNAs in TH. A total of 32978 (19097coding) and 27905 (17911 coding) transcripts were detected with at least one read in TH and SYN, respectively. We also measured protein abundances by mass-spectrometry based proteomics and Data Independent Acquisition (DIA). A total of 4083 and 3221 proteins were detected with non-null peptide absolute intensity in TH and SYN, respectively. The percentage of proteins for which also the corresponding transcript was detected is 77% in TH (N = 3176) and 98% in SYN (N = 3168).

Principal Component Analysis (PCA) (**Figure S1I**) visualized three levels of sample separation based on global transcript and protein expression. TH and SYN samples are separated on PC 1 (89% and 52% of variance explained, respectively, for transcripts and proteins), demonstrating that subcellular origin of samples is the major source of expression variation. The lower separation of TH and SYN at protein level could represent a biological phenomenon suggesting a more homogeneous protein composition across the ages or could be due to the much lower depth of the data generated by proteomics as opposed to transcriptomics. The respective enrichment in SYN of synaptic proteins and transcripts and depletion of nuclear proteins and transcripts for each of the three age steps was further confirmed by computing the probability distribution functions of the enrichment scores and visualizing them as density plots for these two gene- and protein-sets (**Figure S1J-K**). We further assessed the quality of our SYN preparation by comparing our transcriptomic- and proteomic-data with three public synaptosomal datasets and detected significant positive correlation in all three comparisons (Ouwenga et al., 2017; You et al., 2015; Zappulo et al., 2017)(**Table S2** and **Figure S2**).

### Ribosomal proteins are translationally repressed in synaptosomes

We selected genes whose respective transcript and protein level are differentially expressed in either direction between SYN and TH in adult animals (adjusted p-value>0.05, Fisher’s meta-analysis) (Lury & Fisher, 1972). This selection left 2717 pairs of matched proteins and transcripts. Synaptosomal enrichment/depletion at both protein and transcript level for these genes is reported as scatter plot in **Figure 1A**. The first quadrant of this plot contains genes whose transcript- and protein-product show synaptic enrichment; notably, this quadrant shows an enrichment of genes coding for synaptic proteins, as shown also in **Figure 1D**. The third quadrant contains genes that show synaptic depletion of both the respective transcript and protein, the overrepresented GO categories within this gene set show the expected relation to nuclear proteins (**Figure 1D**). The second and the fourth quadrants contain genes whose corresponding proteins and transcripts show opposite directions of enrichment. The second quadrant contains synaptically-depleted proteins whose transcripts are enriched in the synaptosome. The overrepresented GO categories of this gene set correspond mainly to organelles such as the mitochondrion and the ribosome, in line with previous studies (**Figure 1D**). These likely correspond to transcripts that are translationally repressed or proteins whose half-life is shorter in the synapses. The fourth quadrant contains proteins enriched in SYN whose transcripts are more abundant in TH. They are enriched for terms related to vescicle components that are involved in soma-processes trafficking or in synaptic vescicle trafficking (**Figure 1D**). The proteins found in this quadrant are possibly synthesized at somatic level and transported to the neuronal processes. In fact, it has been extensively reported that synaptic vesicle (SV) proteins are synthesized at somatic level and transported to the presynaptic terminal along the axon (Ahmari et al., 2000; Tao-Cheng, 2020; Watson et al., 2023)To confirm that transcripts in the second quadrant of **Figure 1A** (i.e., transcript enriched and protein depleted) are translationally repressed, we performed ribosomes pull-down both from SYN and TH, using a Sucrose Cushion (SC). We then isolated ribosome-associated RNA and performed RNA-seq after rRNA depletion, obtaining an average number of reads of 39.6±3.6 millions. We then calculated the translational efficiency (TE) dividing RPKMs of ribosomal-associated RNAs in SYN (SC_SYN) by RPKMs of ribosomal-associated RNAs in TH (SC_TH). The probability distribution functions of TE of genes belonging to the first and second quadrant of Figure 1A were estimated by Gaussian kernel density and are reported as density plots in **Figure 1B**. We found that genes of the second quadrant are significantly less associated to ribosomes as compared to genes of the first quadrant (p value = 2.546*10^-06^, estimated by Wilcoxon Test). We also analyzed specifically TE of synaptically enriched genes belonging to the GO categories ribosomes (GO:0005840), respiratory chain (GO:0070469) and synapse (GO:0045202) (**Figure 1C**). It is apparent that translation is higher for synaptic transcripts, which are all actively translated at synaptic level as compared to genes belonging to ribosome and respiratory chain categories.

**Figure 1.**
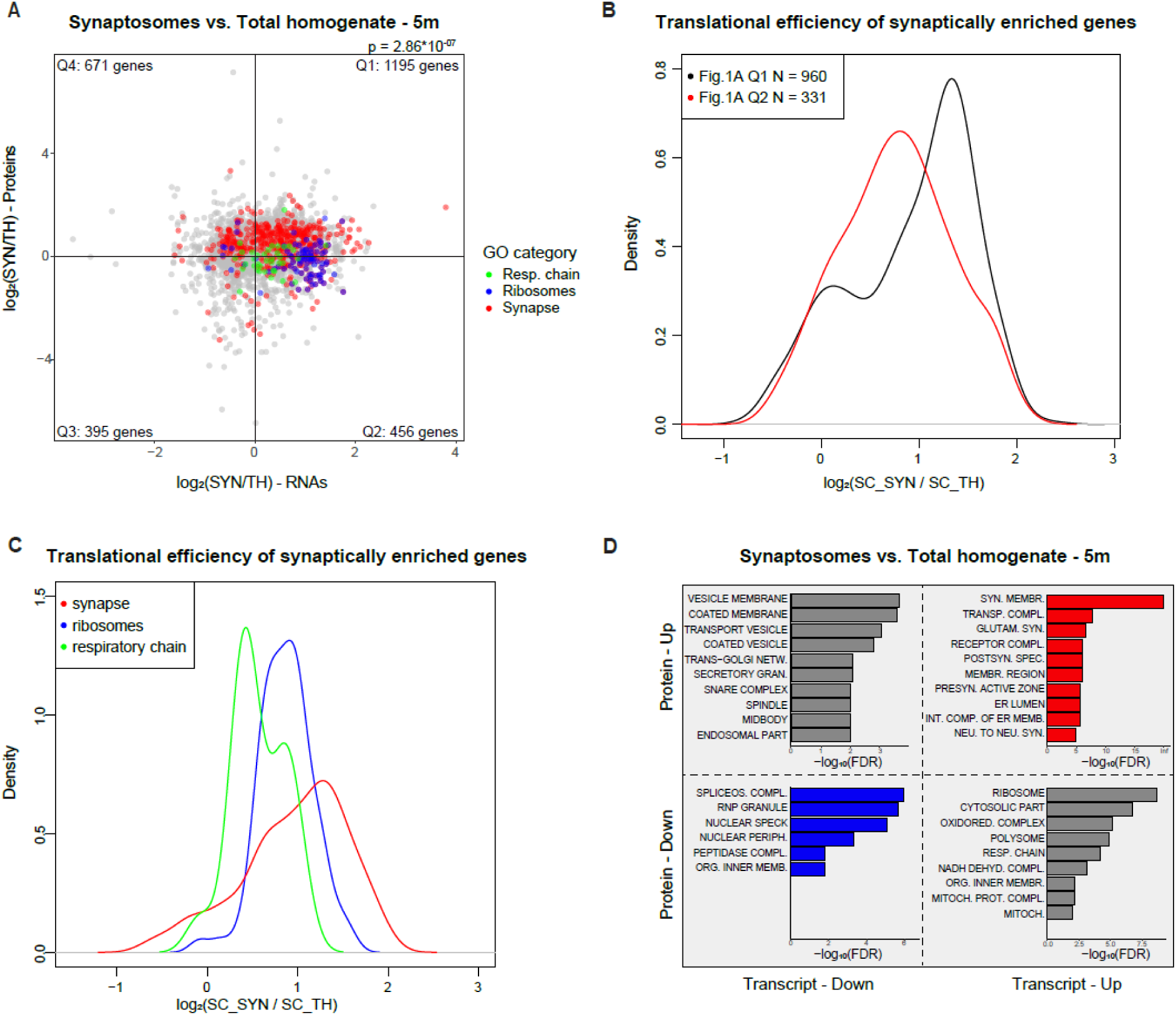
Synaptosomes are enriched in transcripts coding for synaptic proteins. **A)** Proteins and RNAs enrichment are plotted as log of the ratio synaptosome vs. total homogenate. Quadrants are named clockwise, starting from the upper right one. The distribution of genes in each quadrant is not uniform according to Fisher’s exact test (p = 2.86*10^-07^). Protein and RNA fold changes are correlated using Spearman’s correlation (rho = 0.141, p = 1.342*10^-13^). Genes belonging to specific Gene Ontology (GO) terms are highlighted in different colors, red = synapses, blue = ribosome, green = respiratory chain as indicated also in the legend. TH = Total homogenate, SYN = Synaptosomes. **B)** The ratio between ribosome-associated transcripts (translatome) in SYN and TH of genes of quadrant I and II of panel A is visualized in form of density plot. The means are significantly different according to Wilcoxon Test (p value = 2.546*10^-06^). **C)** Probability density plot of the translational efficiency of genes in panel A. Genes belonging to specific Gene Ontology (GO) terms are highlighted in different colors, red = synapses, blue = ribosome, green = respiratory chain as indicated also in the legend. Differences in means are significant according to Kruskal-Wallis Test (p = 0.0001). **D)** Top10 GO categories significantly enriched in each of the 4 quadrants of panel A. Overrepresentation analysis (ORA) was performed with WebGestalt.

We repeated the comparison between proteome and transcriptome of SYN and TH using the same procedure used for the other two time points, yielding, respectively, 3142 and 3062 pairs of matched proteins and transcripts in adult and old animals (**Figure S3A,E**). No substantial change in the genes enriched at synaptic level was observed during aging (**Figure S3D,H**). The most relevant changes in GO categories affect genes that are enriched at transcript but not protein level in synaptosomes (located in the second quadrant). Although these genes appear to have a lower TE compared to those located in Q1 also in young and old animals (**Figure S3B,F**), GO categories related to ribosomes and respiratory chain are overrepresented in the second quadrant in young and adult animals, but not in old animals. Notably, respiratory chain GO category is overrepresented in Q3 (**Figure S3D,H**) at 18 months, indicating that mitochondrial respiration in old animals is impaired both at somatic and synaptic level.

### Non-coding RNAs are enriched in synaptosomes

Some non-coding RNAs (ncRNAs) are reported to be particularly enriched in synapses (Rybak-Wolf et al., 2014; Chen et al., 2017). In our dataset, we detected 26431 genes differentially expressed between SYN and TH, 35% of which are annotated as ncRNAs (9397 genes). In particular, 10871 transcripts, out of which 827 are annotated as ncRNAs, were significantly enriched in SYN. We compared these genes with synaptically-localized ncRNAs extracted from three different datasets (You et al., 2015; Ouwenga et al., 2017; Zappulo et al., 2017) and found 51 ncRNA significantly enriched in synapses in our data and You *et al*. data (**Figure S4A**) and 289 non-coding genes significantly enriched in synaptic compartment in both our dataset and Zappulo *et al*. dataset (**Figure S4C**). As expected, we only found 9 ncRNAs enriched in SYN both in our dataset and Ouwenga *et al*. TRAP-seq data (**Figure S4B**), as this dataset is particularly enriched of ribosome-bound transcripts. We confirmed that ncRNAs significantly enriched in SYN (N = 827) are translationally repressed by comparing them with Zappulo et al. Ribo-seq data, using as control protein coding RNAs that are significantly enriched in SYN (N = 4083). Probability distribution functions of translation rates were estimated by Gaussian kernel density and are reported as density plots in **Figure S4D**. Synaptic enrichment of some long ncRNA was further validated by qPCR (**Figure S4E-G**).

### RNA-protein decoupling in aging synaptosomes

To gain insights on the relationship between transcript and protein abundance, we performed Spearman’s correlation between RNA Reads Per Kilobase Million (RPKMs) and the corresponding protein absolute intensities for each time point separately for TH and SYN. We detected a significant higher correlation in total homogenate as opposed to synaptosomes (p=0.0002 evaluated by two-way ANOVA), likely due to transport of some proteins from the soma. We also observed a decrease in correlation as a function of age (p < 0.0001 evaluated by two-way ANOVA) (**Figure 2A**) indicating age-dependent decoupling between protein and transcript, as previously observed in total brain homogenates of primates and killifish (Kelmer Sacramento et al., 2020; Wei et al., 2015). To analyze transcript- and protein-regulation during aging in TH and SYN, we computed Spearman’s correlation of RPKMs or protein absolute intensities with age for each gene. Rho coefficients were analyzed using Generally Applicable Gene Enrichment (GAGE) (Luo et al., 2009a) to obtain Gene Ontology categories enriched for genes whose expression is positively- or negatively correlated with age (**Figure 2B-C** and **Figure S5A** for TH; **Figure 2D-E** and **Figure S5B** for SYN). In TH, transcripts and proteins related to spliceosomal complex and ribosomes display an opposite behavior with aging. At transcript level, in fact, these genes increase with age, while they decrease with age at protein level. The same phenomenon of protein-transcript decoupling was observed also in the turquoise killifish *Nothobranchius furzeri* (Sacramento et al., 2019). Some examples are depicted in **Figure S5C-D**.

**Figure 2.**
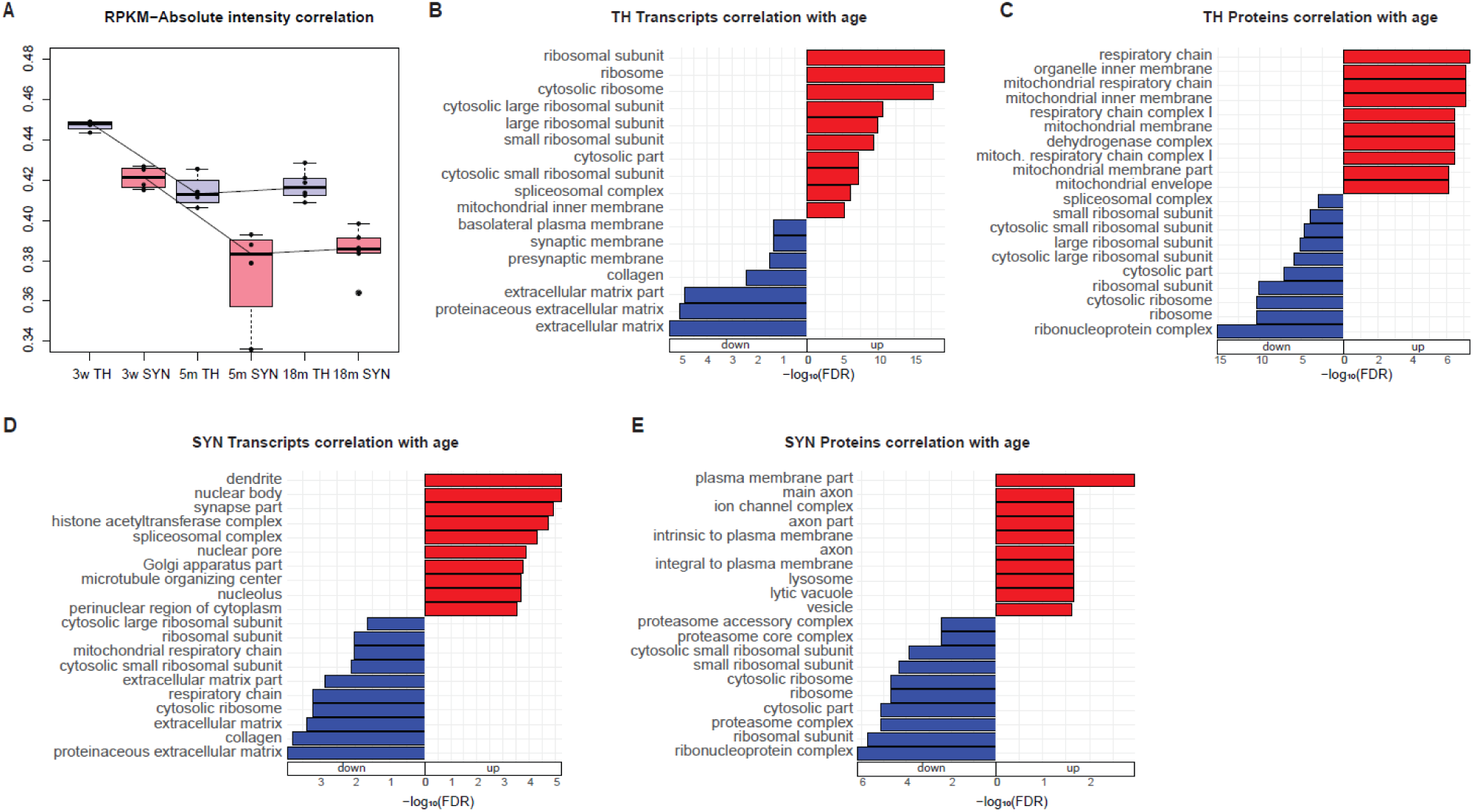
Age-dependent regulation of protein and transcript. **A)** Box plot of the proteome-wide correlation between transcript RPKMs and protein absolute intensities in total homogenate (TH) samples, colored in blue and synaptosomal (SYN) samples, colored in red, at 3 weeks, 5 months and 18 months. Differences between conditions were evaluated by two-way ANOVA. **B)** Generally Applicable Gene Enrichment (GAGE) of the correlation of TH transcripts with age. Only the top10 Cellular Component terms are shown. **C)** GAGE of the correlation of TH proteins with age. Only the top10 Cellular Component terms are shown. **D)** GAGE of the correlation of SYN transcripts with age. Only the top10 Cellular Component terms are shown. **E)** GAGE of the correlation of SYN proteins with age. Only the top10 Cellular Component terms are shown.

We also examined the correlation between age-dependent regulation in SYN and TH during aging, both at transcriptomic and proteomic level. The comparison of transcripts regulated between young and adult stage in SYN and TH yielded 2878 genes significantly regulated in either of the two compartments. The Fold Change of these genes in total homogenate and synaptosomes is highly correlated (rho = 0.811 p value < 2.2*10^-16^ calculated with Spearman’s correlation), suggesting that transcript changes in the synapse are driven by the soma (**Figure 3A**). The same process was repeated for transcripts regulated during aging, yielding 4364 genes commonly regulated in the two compartments. During aging, the correlation between TH and SYN drastically decreases (rho = 0.038, p value = 0.011, calculated with Spearman’s correlation), suggesting that synaptic compartment becomes decoupled from the soma (**Figure 3B**). We observed the same phenomenon at proteome level, with a correlation of the two compartments decreasing from 0.615 (p value < 2.2*10^-16^) during adulthood (**Figure 3C**) to 0.259 (p value = 4.174*10^-16^) during aging (**Figure 3D**), and at the level of the translatome, with a correlation of the two compartments decreasing from 0.713 (p value < 2.2*10^-16^) during adulthood (**Figure 3E**) to 0.146 (p value = 6.024*10^-09^) during aging (**Figure 3F**). While genes coding for synaptic proteins are, as expected, stably expressed and translated during development both in TH and SYN (**Figure S6A,C,E**), the genes more affected from the decoupling are those coding for ribosomal proteins. More specifically, their transcripts increase and their proteins decrease in TH while the opposite is observed in SYN (**Figure S6B** and **Figure S6D**, respectively).

**Figure 3.**
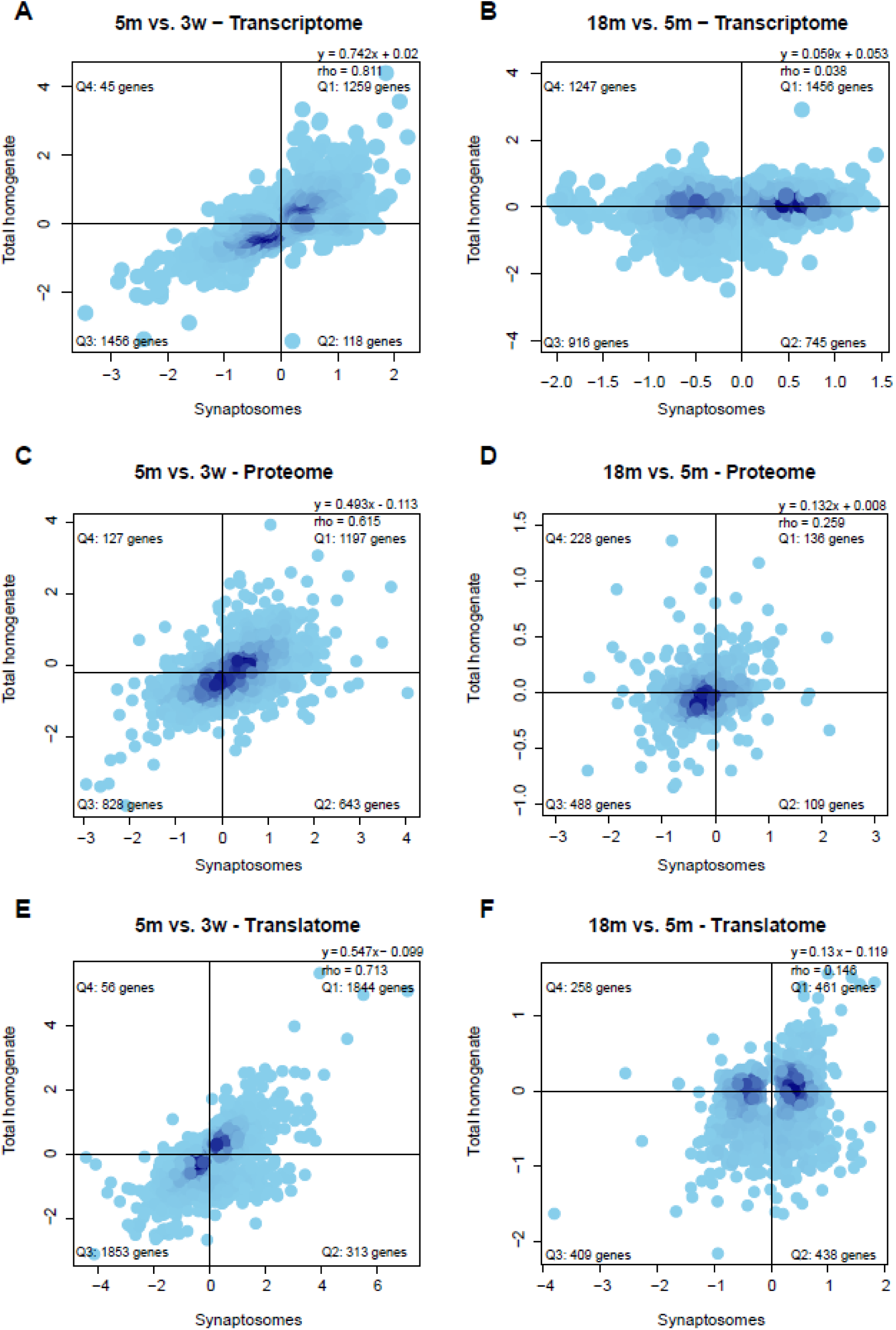
Soma-synapse decoupling during aging. **A)** Transcript regulation plotted as log of the ratio 5m vs. 3w in TH and SYN. Quadrants are named clockwise, starting from the upper right one. The distribution of genes in each quadrant is not uniform according to Fisher’s exact test (p < 2.2*10^-16^). SYN and TH fold changes are correlated based on Spearman’s correlation (rho = 0.811, p < 2.2*10^-16^). **B)** Transcript regulation plotted as log of the ratio 18m vs. 5m in TH and SYN. Quadrants are named clockwise, starting from the upper right one. The distribution of genes in each quadrant is not uniform according to Fisher’s exact test (p = 7.856*10^-09^). SYN and TH fold changes are weakly correlated using Spearman’s correlation (rho = 0.038, p = 0.011). **C)** Protein regulation plotted as log of the ratio 5m vs. 3w in TH and SYN. Quadrants are named clockwise, starting from the upper right one. The distribution of genes in each quadrant is not uniform according to Fisher’s exact test (p < 2.2*10^-16^). SYN and TH fold changes are correlated using Spearman’s correlation (rho = 0.615, p < 2.2*10^-16^). **D)** Protein regulation plotted as log of the ratio 18m vs. 5m in TH and SYN. Quadrants are named clockwise, starting from the upper right one. The distribution of genes in each quadrant is not uniform according to Fisher’s exact test (p = 9.909*10^-11^). SYN and TH fold changes are correlated using Spearman’s correlation (rho = 0.259, p = 4.174*10^-16^). **E)** Ribosome-associated transcript regulation plotted as log of the ratio 5m vs. 3w in TH and SYN. Quadrants are named clockwise, starting from the upper right one. The distribution of genes in each quadrant is not uniform according to Fisher’s exact test (p < 2.2*10^-16^). SYN and TH fold changes are correlated using Spearman’s correlation (rho = 0.713, p < 2.2*10^-16^). **F)** Ribosome-associated RNAs transcript regulation plotted as log of the ratio 18m vs. 5m in TH and SYN. Quadrants are named clockwise, starting from the upper right one. The distribution of genes in each quadrant is not uniform according to Fisher’s exact test (p = 8.305*10^-07^). SYN and TH fold changes are correlated using Spearman’s correlation (rho = 0.146, p = 6.24*10^-09^).

To explore further the age-dependent transcriptome/proteome decoupling, we analyzed the correlation between RNA and proteins during development and aging, both in TH (**Figure 4A,D** and **S7A,B**) and SYN (**Figure 4G,J** and **S7C,D**), highlighting in the scatterplots the GO categories that are most affected by aging, namely respiratory chain and ribosome. This correlation decreases from 0.221 (p value < 2.2*10^-16^) to −0.011 (p value = 0.743) in TH and from 0.236 (p value < 2.2*10^-16^) to 0.028 (p value = 0.246) in SYN. We thus calculated a decoupling score for each gene defined as the ratio between protein and transcript changes: positive decoupling indicates that the protein increases more (for is reduced less) than expected from transcript regulation and negative decoupling that the protein increases less (or decreases more) than expected from transcript regulation (Fraia et al., 2023) and used decoupling scores of each gene as input for GAGE (**Figure 4B,E,H,K**). This analysis showed that ribosomal genes have a negative decoupling during development and aging in TH. In SYN, on the other hand, ribosomal genes show a negative decoupling during development and a positive decoupling during aging. Analysis of the correlation of decoupling with changes in translational efficiency (TE) in old SYN confirmed that positive decoupling correlates with an increase in TE and that higher protein concentration is the result of higher rates of protein synthesis (**Figure 4L**).

**Figure 4.**
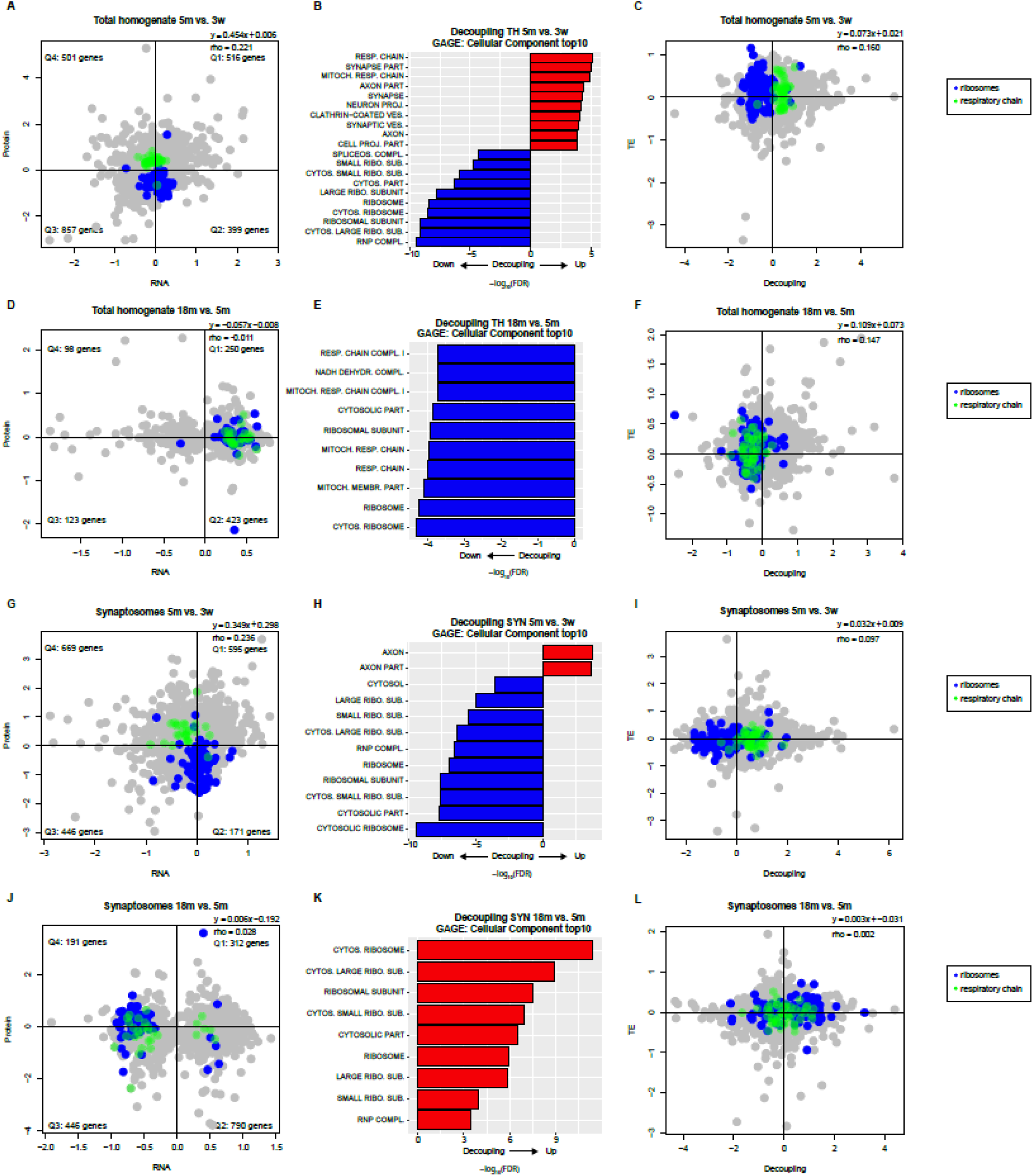
Transcriptome/proteome decoupling with aging in TH and SYN. **A)** Proteins and RNAs enrichment in TH are plotted as log of the ratio 5m vs. 3w. Quadrants are named clockwise, starting from the upper right one. The distribution of genes in each quadrant is not uniform according to Fisher’s exact test (p < 2.2*10^-16^). Protein and RNA fold changes were correlated using Spearman’s correlation (rho = 0.221, p < 2.2*10^-16^). Genes belonging to specific Gene Ontology (GO) terms are highlighted in different colors, blue = ribosome, green = respiratory chain as indicated also in the legend. **B)** Generally Applicable Gene Enrichment (GAGE) of the decoupling of transcripts and proteins in TH during development. Positive decoupling indicates higher protein level than expected from transcript regulation and negative decoupling less protein level than expected from transcript regulation. Only the top10 categories ranked on log2 of the p-value are shown. **C)** Decoupling and Translational Efficiency (TE) in TH are plotted as log of the ratio 5m vs. 3w. Decoupling and TE fold changes were correlated using Spearman’s correlation (rho = 0.160, p < 2.2*10^-16^). Genes belonging to specific Gene Ontology (GO) terms are highlighted in different colors, blue = ribosome, green = respiratory chain as indicated also in the legend. **D)** Proteins and RNAs enrichment in TH are plotted as log of the ratio 18m vs. 5m. Quadrants are named clockwise, starting from the upper right one. The distribution of genes in each quadrant is uniform according to Fisher’s exact test (p = 0.0673). Protein and RNA fold changes were not significantly correlated according to Spearman’s correlation (rho = −0.011, p = 0.743). Genes belonging to specific Gene Ontology (GO) terms are highlighted in different colors, blue = ribosome, green = respiratory chain as indicated also in the legend. **E)** GAGE of the decoupling of transcripts and proteins in TH during aging. Only the top10 categories are shown. **F)** Decoupling and TE in TH are plotted as log of the ratio 18m vs. 5m. Decoupling and TE fold changes were correlated using Spearman’s correlation (rho = 0.147, p < 2.2*10^-16^). Genes belonging to specific Gene Ontology (GO) terms are highlighted in different colors, blue = ribosome, green = respiratory chain as indicated also in the legend. **G)** Proteins and RNAs enrichment in SYN are plotted as log of the ratio 5m vs. 3w. Quadrants are named clockwise, starting from the upper right one. The distribution of genes in each quadrant is not uniform according to Fisher’s exact test (p = 6.165*10^-16^). Protein and RNA fold changes were correlated using Spearman’s correlation (rho = 0.236, p < 2.2*10^-16^). Genes belonging to specific Gene Ontology (GO) terms are highlighted in different colors, blue = ribosome, green = respiratory chain as indicated also in the legend. **H)** GAGE of the decoupling of transcripts and proteins in SYN during development. Only the top10 categories are shown. **I)** Decoupling and TE in SYN are plotted as log of the ratio 5m vs. 3w. Decoupling and TE fold changes were correlated using Spearman’s correlation (rho = 0.097, p = 2.515*10^-09^). Genes belonging to specific Gene Ontology (GO) terms are highlighted in different colors, blue = ribosome, green = respiratory chain as indicated also in the legend. **J)** Proteins and RNAs enrichment in SYN are plotted as log of the ratio 18m vs. 5m. Quadrants are named clockwise, starting from the upper right one. The distribution of genes in each quadrant is uniform according to Fisher’s exact test (p = 0.475). Protein and RNA fold changes were correlated using Spearman’s correlation (rho = 0.028, p = 0.247). Genes belonging to specific Gene Ontology (GO) terms are highlighted in different colors, blue = ribosome, green = respiratory chain as indicated also in the legend. **K)** GAGE of the decoupling of transcripts and proteins in SYN during aging. Only the top10 categories are shown. **L)** Decoupling and TE in SYN are plotted as log of the ratio 18m vs. 5m. Decoupling and TE fold changes were correlated using Spearman’s correlation (rho = 0.002, p = 0.875). Genes belonging to specific Gene Ontology (GO) terms are highlighted in different colors, blue = ribosome, green = respiratory chain as indicated also in the legend.

We confirmed the increase of ribosomal proteins in SYN during aging by a Western blot with an independent group of samples (**Figure 5A-B**, p = 0.00717 evaluated by two-way ANOVA). Similarly, the decrease od transcripts coding for ribosomal proteins was confirmed by means of qPCR (**Figure 5C**, p = 0.00154 evaluated by two-way ANOVA). Moreover, we evaluated the amount of ribosomal RNA (rRNA) with a bioanalyzer (**Figure 5D**). This datum indicates that higher ribosomal proteins correlate with an increase in fully-assembled ribosome at synaptic level. Finally, a staining of a ribosomal protein and RNA on isolated synaptosomes (Richter et al., 2018), further corroborated our data (**Figure 5E**, p<0.0001 evaluated by Kruskal-Wallis test).

**Figure 5.**
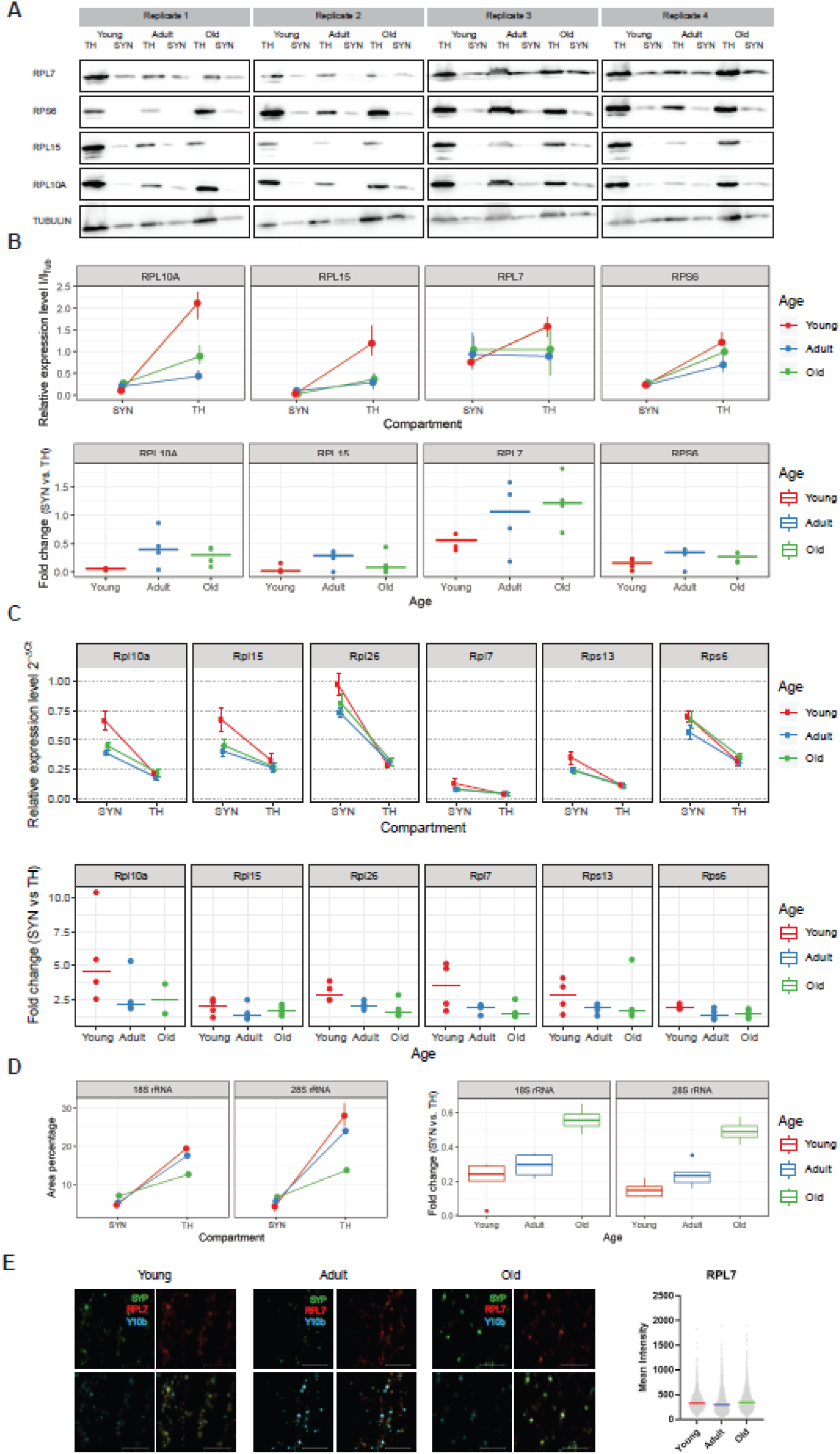
Validation of ribosomal transcripts- and proteins-regulation in synaptosomes during aging. **A)** Western blot of ribosomal proteins using total cortical homogenate and synaptosomes of young, adult and old animals. Tubulin was used as loading control in all the experiments. **B)** Upper panel: mean of protein intensity quantification in synaptosomes (SYN) and total homogenate (TH) relative to tubulin for each time point of panel A. Lower panel: Fold change of ribosomal proteins in SYN normalized to TH fold changes. Red indicates young animals, blue adult animals and green old animals. **C)** Upper panel: mean of ribosomal transcripts expression in synaptosomes (SYN) and total homogenate (TH) relative to Ldhb for each time point using independent samples. Lower panel: Fold change of ribosomal transcripts in SYN normalized to TH fold changes. Red indicates young animals (N = 4), blue adult animals (N = 4) and green old animals (N = 4). **D)** Ribosomal RNA (rRNA) in Total Homogenate (TH) and Synaptosomes (SYN). Box plot of the fold change of 18S and 28S between SYN and TH across the ages. Red indicates young animals (N = 4), blue adult animals (N = 4) and green old animals (N = 4). **E)** Immunofluorescence of SYN isolated from young, adult and old animals (N = 4 for each time point). Synaptic protein synaptophysin (SYP) stained in green, in red Ribosomal protein RPL7 and in blue Ribosomal RNA. Bar 5µm. Right panel: quantification of mean fluorescence intensity of RPL7 using SYP as reference. Each dot represents a single synaptosome, bars indicate the mean with 95% Confidence Interval (CI).

Taken together, the higher levels of ribosomal proteins, correlated with an increase in translational efficiency (TE) in aging synaptosomes, pinpoint to a compensatory mechanism that enhances protein synthesis at the synapse, possibly to counteract the decline in overall protein homeostasis.

### Differential expression of transcript isoforms in aging synaptosomes

In order to identify transcript isoforms that differentially expressed in SYN with aging, we performed paired-end long read sequencing on TH and SYN from young, adult and old mice of ages corresponding to the previous experiments. We applied both DIEGO (Doose et al., 2018) and LeafCutter (Y. I. Li et al., 2018). In both cases, Differentially Expressed Junctions (DEJs) were identified for each of the age comparisons SYN 3w/5m, SYN 3w/18m and SYN 5m/18m. Both software identify transcript isoforms by determining differentially expressed junctions based exclusively on split reads information and do not make use of a priori gene models, like DEXSeq (Anders et al., 2012). The two software, however, use different methods for the detection of statistically significant DEJs. DIEGO analyzes each splice junction separately and assigns a q-value to its differential usage. LeafCutter analyzes clusters, i.e. a set of junctions connected to each other through common acceptor or donor splice sites and assigns an adjusted p-value to each cluster based on differential connectivity within the cluster. In the three aforementioned age comparisons 3099, 1496 and 2192 DEJs (DIEGO) and 4249, 4175 and 4036 clusters (LeafCutter) were identified, respectively. Both software identified only a negligible number of DEJs that could not be mapped to an annotated gene or that spanned an unrealistically large range (>106bp). For each age comparison, we evaluated the percentage of DEJs with either: i) both donor and acceptor splice sites annotated, ii) only one of the two splice sites annotated and iii) both splice sites unannotated (**Figure S8A-B**). LeafCutter identified more junctions with at least one unannotated splice site as compared to DIEGO (18.74% in LeafCutter and 8.95% in DIEGO, on average). Next, we evaluated the percentage of DIEGO-derived DEJs (**Figure S8C**) and LeafCutter-derived clusters (**Figure S8D**) mapping on the 5’UTR, on the coding sequence (CDS), on the 3’UTR and on the introns. Most splice sites map to the UTRs and the CDS; however, we detected a higher percentage of LeafCutter-identified splice sites mapping on the 5’UTR and on introns, suggesting either that LeafCutter identifies more unannotated exons and 5’UTR isoforms, or that it has a higher rate of false positive calls, as compared to DIEGO. We performed GO terms overrepresentation analysis on the top 1000 q-value ranked DIEGO-derived DEJs (**Figure S8E-G**) and on the top 1000 adjusted p-value ranked LeafCutter-derived clusters (**Figure 6A-C**), to gain insights into the function of age-dependent differentially spliced synaptic transcripts. The GO category GTPase binding (GO:0051020) resulted preeminent in all three comparisons. GTPase activity is involved in a multitude of processes that are important in synaptic function, like signal transduction at the level of G protein-coupled receptors, neurotransmitter release, protein synthesis and synaptic plasticity. This enzymatic activity is regulated by various GTPase binding proteins, like guanine nucleotide exchange factors (GEFs), which catalyze the exchange of GDP with GTP and the reactivation of the GTPase, GTPase activating proteins (GAPs), which increase the GTPase activity, and guanosine nucleotide dissociation inhibitors (GDIs), that inhibit the exchange of GDP with GTP and therefore maintain the GTPase in its inactive state.

**Figure 6.**
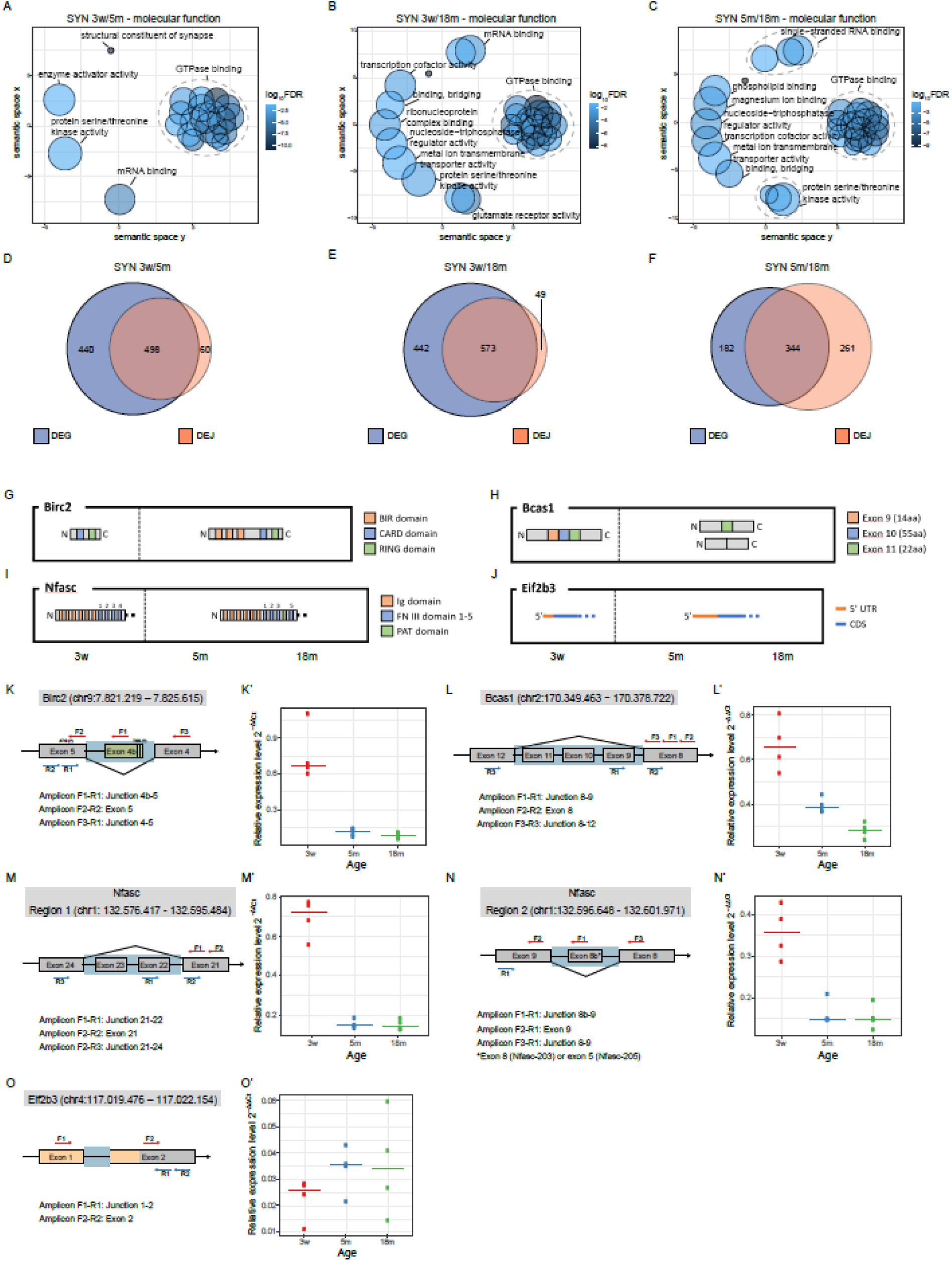
Differentially spliced genes (LeafCutter) show over-representation of GTPase binding activity and significantly overlap with differentially expressed genes (DEGs) in the same age comparison. **A-C)** Gene Ontology (GO) categories (Molecular Function) that are overrepresented in the list of genes to which the 1000 most significant clusters (ranked on adjusted p-value) map in the three age comparisons SYN 3m vs. 5m (**A**), SYN 3w vs. 18m (**B**) and SYN 5m vs. 18m (**C**) are represented by REVIGO through multidimensional scaling using a measure of semantic dissimilarity (the distances between dots, that represent single GO categories, are inversely related to the similarity of the categories). Ellipses delimit manually identified clusters of highly similar GO categories, and a representative term of each cluster is indicated. **D-F)** Overlap of the list of GO categories that are overrepresented in the list of genes to which the 1000 most significant clusters (lowest adjusted p-value) map (orange) and the list of genes that are differentially expressed (adjusted p-value < 0.01) (blue) in the three age comparisons SYN 3m vs. 5m (**D**), SYN 3w vs. 18m (**E**) and SYN 5m vs. 18m (**F**). The overlap is highly significant at all three age comparisons but smaller for the SYN 5m vs. 18m comparison, given that both the comparisons 60/498 vs. 261/344 and 49/573 vs. 261/344 are significantly different (Fisher’s exact test, p-value < 0.0001). **G-J)** Schematic representation of the predominant protein and transcript isoforms of the three genes *Birc2* (**G**), *Bcas1* (**H**), *Nfasc* (**I**) and *Eif2b3* (**J**) in young, adult and old synapses. The sizes and distances of the different domains are not to scale. **K-O’)** Experimental validation of candidate junction. **K)** Schematic representation of the primer pairs used for RT-qPCR and RT-PCR validations for *Birc2*. Please note that the coding sequence of *Birc2* is annotated on the – strand. Exons are represented as grey boxes and introns as black lines (not to scale). The green box represents the unannotated exon 4b. A light blue background represents the region spanned by the identified DEJ. Putative TSSs and ATG sites are shown, respectively, on exon 4b and 5. The dotted lines at the 3’ end of exon 4b indicate that this is not clearly defined. **K’)** Validation by RT-qPCR. The relative expression of the Junction 4b-5 is reported as normalized by expression of the Exon 5. **L)** Schematic representation of the primer pairs used for RT-qPCR and RT-PCR based validation for *Bcas1*. Please note that the coding sequence of *Bcas1* is annotated on the – strand. Exons are represented as grey boxes and introns as black lines (not to scale). A light blue background represents the region spanned by the identified DEJ. **L’)** Validation by RT-qPCR. The relative expression of the Junction 8-9 is reported as normalized by expression of the Exon 8. **M,N)** Schematic representation of the primer pairs used for RT-qPCR and RT-PCR based validation of Region 1 and Region 2 of *Nfasc*. Please note that the coding sequence of *Nfasc* is annotated on the – strand. Exons are represented as grey boxes and introns as black lines (not to scale). Light blue backgrounds represent the two regions. **M’,N’)** Validation by RT-qPCR. The relative expression of the Junctions 21-22 (**M’**) and 8b-9 (**N’**) are reported as normalized by expression of Exons 21 (**M’**) and 9 (**N’**). **O)** Schematic representation of the primer pairs used for RT-qPCR based validation for *Eif2b3*. Exons are represented as grey boxes and introns as black lines (not to scale). Exons or parts of exons that are part of the 5’UTR are represented in orange. A light blue background represents the region spanned by the identified DEJ. **O’)** Validation by RT-qPCR. The relative expression of the Junction 1-2 is reported as normalized by expression of the Exon 2.

Furthermore, we compared the overlap between the GO categories enriched in the DEGs (adjusted p-value <0.01) and in the genes that contain DEJs identified both with DIEGO (top 1000 q-value ranked DEJs) and LeafCutter (top 1000 adjusted p-value ranked clusters) (**Figure S8H-J** and **Figure 6D-F**). We observed a larger overlap between DEGs and DEJs in the contrast SYN 3w vs 5m as compared to the contrast SYN 5m vs. 18m, with only 10.7% (60/558) DEJs not DEG (**Figure 6D**) as opposed to the contrast SYN 5m vs. 18m with 43% (261/605) DEJs not DEG (**Figure 6F**, p<10-6, Fisher’s exact test), suggesting that during aging regulation of transcript abundance and alternative splicing become more decoupled. As a general rule, changes in synaptic transcriptome during aging, might be imputable either to changes in relative expression of certain transcripts (DEGs) or to a transcript isoform switch (DEJs), where the two processes can occur independently. We observed a significant overlap in GO categories enriched in DEGs and genes containing DEJs when comparing SYN of young animals with those of adult and old animals, suggesting that, during development, the same biological processes are modulated via changes in both transcript abundance and transcript isoform. On the other hand, during aging, the two phenomena diverge, indicating that synaptic gene expression might be regulated independently by changes in transcript abundance and changes in transcript isoform.

We selected a subset of DEJs which were preeminent in at least one of the three age comparisons, according to either DIEGO or LeafCutter (**Table 1**) and examined the variation in transcript abundance in the three age comparisons (**Table 2**). Bcas1 and Nfasc showed a marked decrease upon aging, while the other genes were not differentially expressed in the analyzed time points. A summary of the age-dependent protein and transcript isoform changes is shown in **Figure 6G-J**. The change in the usage of the selected junctions was validated through RT-qPCR (**Figure 6K-O’**) and RT-PCR (**Figure S9A-C**) using an independent set of samples. In all the considered genes, the junction is differentially used due to differential expression of adjacent exons or of exons spanning that junction, we therefore designed three primer pairs, one amplifying a “normalizer exon”, immediately adjacent to the selected junction, one amplifying the junction between the differentially expressed exon and the normalizer exon and the last one amplifying the entire region in which alternative splicing was detected. For the validation through RT-qPCR, we used the gene *Ldhb* as normalizer, due to its stable expression, between both TH and SYN at the considered time points.

**Table 1.**
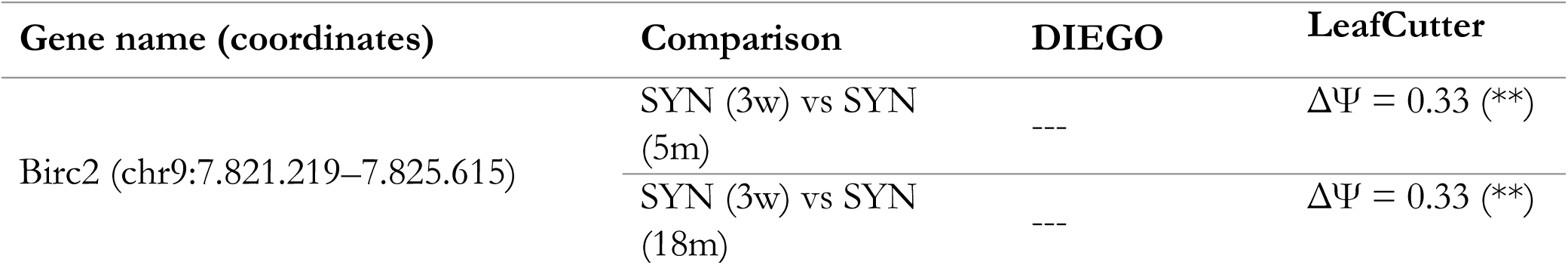
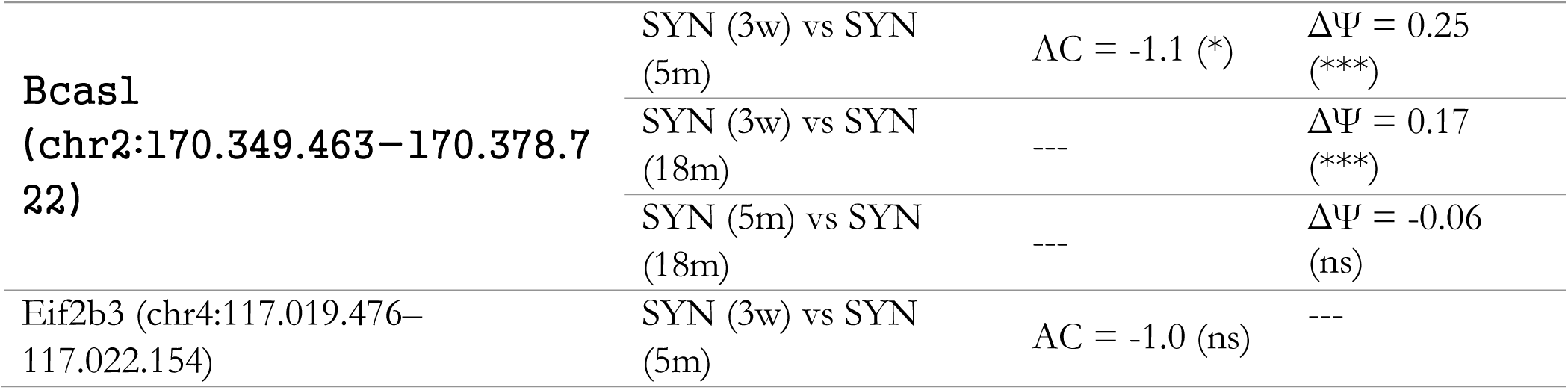
List of candidate junctions chosen for validation. AC: Abundance Change; Significance values of q-value (DIEGO) and adjusted p-value (LeafCutter) are indicated in the following way: ns: > 0.05, *: < 0.05, **: < 0.01, ***: < 0.001. AC values are rounded to the nearest tenth, q-value and adjusted p-value value to the nearest thousandth and Psi to the nearest hundredth.

**Table 2.** Log2 fold change (log2FC) and the corresponding adjusted p-values of the candidate genes in the three age comparisons SYN 3m vs. 5m, SYN 3w vs. 18m and SYN 5m vs. 18m. Adjusted p-values are indicated in the following way: ns: > 0.05, *: < 0.05, **: < 0.01, ***: < 0.001. Log2FC values are rounded to the nearest hundredth.

|  | SYN (3w) vs SYN (5m) | SYN (3w) vs SYN (18m) | SYN (5m) vs SYN (18m) |
| --- | --- | --- | --- |
| **Birc2** | log<sub>2</sub>FC = 0.17 (ns) | log<sub>2</sub>FC = 0.31 (ns) | log<sub>2</sub>FC = 0.13 (ns) |
| **Bcas1** | log<sub>2</sub>FC = 0.87 (***) | log<sub>2</sub>FC = 1.11 (***) | log<sub>2</sub>FC = 0.24 (ns) |
| **Nfasc** | log<sub>2</sub>FC = 1.31 (***) | log<sub>2</sub>FC = 1.20 (***) | log<sub>2</sub>FC = -0.12 (ns) |
| **Eif2b3** | log<sub>2</sub>FC = -0.38 (ns) | log<sub>2</sub>FC = -0.08 (ns) | log<sub>2</sub>FC = 0.29 (ns) |

#### Birc2

BIRC2 is an Inhibitor of Apoptosis Protein (IAP) and inhibits apoptosis by binding Caspases (through BIR domains) and ubiquitinating them (Silke & Meier, 2013). By inspecting the read coverage of Birc2, we identified an unannotated exon (to which we will refer to as “Exon 4b”) in 3-weeks SYN. As shown in **Table S3** and **Figure S9D**, split reads span the junction between exon 4 and exon 5 and the junction between exon 4b and exon 5, but no split read is detectable between exon 4 and 4b; furthermore, the putative unannotated donor splice site at the 3’ end of exon 4b (GTAAGC) is consistent with the consensus sequence “GTAAGT”, thus we hypothesized that an unannotated Transcription Start Site (TSS) might be present at the 5’ end of exon 4b, generating a transcript isoform that does not include exons 1-4 and, consequently, a protein with a shorter N-terminal. In fact, we were able to identify in exon 5 a start codon contained in a Kozak sequence and in frame with the open reading frame (ORF) using ATGpr tool (Salamov et al., 1998). In agreement with these results, the amplicon spanning the junction 4b-5 is more expressed in SYN 3w as compared to SYN 5m and SYN 19m, indicating that the unannotated isoform is only present in young synapses (**Figure 6K-K’** and **Figure S9A**). In the truncated protein isoform, the first 395aa are missing. That means that the three BIR domains are not present in this protein and that only the RING domain (amino acids 565-600) and the CARD domain (amino acids 447-537) remain (**Figure 6G**), where the latter can inhibit the E3 ubiquitin-protein ligase activity by preventing RING domain dimerization (Lopez et al., 2011). We hypothesize that the loss of function of BIRC2 that results from the loss of its BIR domains facilitates the successful activation of the apoptotic signaling in neurons of young animals, which could be important for synaptic pruning and neuronal circuit formation. Alternatively, this truncated protein may act non-canonically and control biological processes other than apoptosis.

#### Bcas1

BCAS1 (Breast Carcinoma Amplified Sequence 1) is a basic protein highly expressed in the brain; its knockout is associated with schizophrenia-like behavioral abnormalities in mice (Fard et al., 2017; Ishimoto et al., 2017). The inspection of the coverage of its transcript (**Figure S9E**) suggests that exons 9-11 are more expressed in SYN 3w as compared to SYN 5m and SYN 18m. In agreement with this result, the amplicon spanning the junction 8-9 is higher expressed in young SYN and progressively decreases its expression with aging (**Figure 6L-L’** and **Figure S9B**). Interestingly, the amplification of the junction 8-12 generates four amplicons, of which only two correspond to the isoforms containing or excluding the exons 9-10-11, while the other two might be further variants containing only some of the three exons (**Figure S9B**). The exons 9, 10 and 11 have all lengths multiple of 3 (respectively 42nt, 165nt and 66nt) and all the possible transcript isoforms are therefore in frame for translation. Therefore, in adult and old synapses the short-length and the mid-length isoforms are more frequent, while in young synapses the long-length isoform is the dominant isoform (**Figure 6H**). Little is known about the structure of the protein and its possible role in synapse physiology. However, various phosphorylation sites are annotated (Huttlin et al., 2010), and some of them are located in the N-terminal region that is coded in isoform 201, but not in isoform 208 transcripts, and in the exons 9-11. The age-dependent translation of the abovementioned transcript isoforms would therefore lead to peptides with different phosphorylation sites, and that, on the other hand, could have an impact on the protein function.

#### Nfasc

Neurofascin is a cell adhesion molecule of the L1 subgroup of the immunoglobulin superfamily that mediates homophilic adhesions through its Ig-like domains. It mediates axon targeting and synapse formation during neural development, and both glial and neuronal isoforms are known. This transmembrane protein is organized in an N-terminal extracellular region, a transmembrane domain and a C-terminal cytoplasmic domain. The first 24aa of the N-terminal represent the signal peptide and are therefore absent in the mature protein. The extracellular domain is composed of six Ig-like domains, five Fibronectin type-III (FN III) domains and one PAT domain, and the various isoforms have different FN III domains and can have or not have the PAT domain (Kriebel et al., 2012; Suzuki et al., 2017). Exon pairs 18-19, 20-21, 22-23, 24-25 and 28-29 (isoform 207) code, respectively, for the FN III domains 1-5, while the exon pair 26-27 codes for the PAT domain.

This transcript is subject to extensive differential splicing. We identified 18 and 20 DEJs with DIEGO and LeafCutter, respectively, in the three age comparisons, which, if merged by interval overlapping, are organized in 6 regions along the transcript, of which all but region 3 and 5 showed any detectable age-dependent change in coverage. Those regions correspond, respectively, to exons 26-29 (Region 1), exons 22-23 (Region 2), exon 8b (Region 4) and exon 1 (Region 6). The inspection of the coverage suggests that exons 26-29 (Region 1) are less expressed in SYN 3w, as compared to SYN 5m and SYN 18m. Furthermore, the inspection of the coverage of exons 22-23 (Region 2) suggests that they are more expressed in SYN 3w, as compared to SYN 5m and SYN 18m. In line with this result, we confirmed that the amplicon spanning the junction 21-22 is expressed the most in SYN 3w (**Figure 6M-M’**). The amplification of the region spanning junctions 21-24 provided a clear result, since we were able to observe two bands, corresponding to the isoforms with and without exons 22-23, of which only the first one is amplified in SYN 3w (**Figure S9C**). Those results suggest that the predominant transcript isoform in young synapses does not code for the 5^th^ FN III domain and the PAT domain (exons 26-29 less expressed), while the predominant transcript isoform in adult and old synapses does not code for the 3^rd^ FN III domain (exons 22-23 less expressed). This suggests that in young synapses the predominant transcript isoform codes for NF155, while in adult and old synapses the predominant transcript isoform codes for NF186 (**Figure 6I**). In fact, it is known that the isoform NF155 is the glial isoform, which is expressed with the onset of myelination, while the isoform NF186 is predominantly expressed in the adult brain, where it stabilizes axo-axonic synapses and mediates the clustering of Na_v_ channels (Kriebel et al., 2012).

Furthermore, when we inspected the coverage of region 4, we hypothesized that exon 8b (51nt) could be more expressed in SYN 3w, compared to SYN 5m and SYN 19m. In agreement with this result, the amplicon spanning the junction 8b-9 resulted to be more expressed in SYN 3w, both in RT-qPCR and RT-PCR (**Figure 6N-N’** and **Figure S9C**). The amplification of the region spanning the junction 8-9 generates three different amplicons, of which two correspond to the isoforms with and without exon 8b. The first one is slightly more expressed in SYN 3w, while the second one is clearly expressed the most in SYN 3w. The third amplicon (slightly longer than 200bp) might represent an unspecific amplification, given that no third splicing isoform can be observed from the coverage data. Given that this region codes for a part of the portion between the 2^nd^ Ig-like domain (exons 7-8) and the 3^rd^ Ig-like domain (exons 9-10), it seems that the predominant transcript isoform in young synapses codes for an 17aa longer variant of this protein segment.

Finally, we evaluated region 6, noticing that two 5’UTR isoforms could be observed: an Nfasc-202/-203 isoform (286/292nt), that can be observed at all three ages, and an Nfasc-201 isoform (427nt), that can be observed only in young synapses (**Figure S9F**). The biological function of these two different 5’UTRs, and the underlying alternative TSSs, has not been inquired yet.

#### Eif2b3

EIF2B is the GEF that catalyzes the reactivation of eIF2alfa, and EIF2B3 is its third (gamma) subunit. The inspection of the coverage of its transcript suggests that exon 1, which is part of the 5’UTR, is less expressed in SYN 3w, compared to SYN 5m and SYN 18m. In other words, in young synapses the isoform 201 (short 5’UTR) seems to be preferred (**Figure 6J**). In agreement with this result, the amplicon spanning the junction 1-2 is less expressed in SYN 3w, compared to SYN 5m and SYN 18m (**Figure 6O-O’**). The isoform 201, which is predominant in young animals, is characterized by a much smaller 5’UTR. Although this isoform is already annotated, its biological function has not been studied yet. The presence of a short 5’UTR can have a variety of implications for the transcript localization, stability and translation rate, and that, on the other hand, could allow for an age-dependent global regulation of synaptic translation.

## CONCLUSION

During adult life, protein and transcript regulation at the synapse are coupled to changes at the level of neuronal cell bodies. During aging, we observe both protein-transcript uncoupling and soma-synapse uncoupling for both proteins and transcripts as well as synapse specific changes in splicing. Aging is associated with synapse-specific translational remodeling that is particularly striking for what concerns transcripts coding for ribosomal proteins. These are enriched but translationally inhibited in young synapses. During aging these transcripts are down-regulated at the synapse but are more associated to ribosomes and the corresponding proteins increase because their translation is specifically induced at synaptic level during aging.

## MATERIALS AND METHODS

### Mouse maintenance

Tissue from wild-type C57BL/6 mice was obtained from the Leibniz Institute on Aging, Fritz Lipmann Institute Jena and from Janvier facility. Mice were housed in standard cages and fed standard lab chow and water ad libitum. Euthanasia was carried out by cervical dislocation in accordance with the European Council Directive of 22 September 2010 (2010/63/EU). The scientific purposes of the experiments were approved by the local authorities and supervised by the local veterinary (Anzeige N. O_AC_04, O_AC_08, O_AC_10 and O_AC_23-27).

### Synaptosome Preparation and Validation

Synaptosomes were extracted from cerebral cortices of 3 weeks-, 5 months- and 18/19 months-old mice as previously described (Gray & Whittaker, 1962; Moczulska et al., 2014). Briefly, the cortex was dissected and transferred to ice-cold synaptosomal buffer (10 mM HEPES, 1 mM EDTA, 2 mM EGTA, 0.5 mM DTT, 0.32 M sucrose, pH 7) containing protein inhibitors (1 tablet/10 mL, Roche). Samples were homogenized using a pestle (Sigma-Aldrich) with 12 strokes on ice. The homogenate was then centrifuged for 10min at 1.000g at 4°C and the supernatant was centrifuged again for 40min at 10.000g at 4°C. The obtained pellet, that contains the synaptosomes, was then resuspended in ice-cold synaptosomal buffer and layered over a discontinuous sucrose gradient (0.8M/1M/1.18M sucrose). The samples were then centrifuged for 2h at 50.512g at 4°C (Beckman Coulter Optima XPN-80 ultracentrifuge). The interface between 1M and 1.18M was withdrawn (synaptosome sample) and stored at −80°C. Aliquots were taken after the homogenization (Total Homogenate samples), after the first centrifugation (Cytosolic Fraction samples) and after the resuspension of the pellet containing the synaptosomes (Raw Synaptosomes samples).

Synaptosome samples were processed for RNA extraction according to (Smalheiser et al., 2014), and RNA was then extracted from the appositely prepared synaptosome samples and from TH samples through acidic phenol-chloroform extraction according to (Chomczynski, 1993). The synaptosomes were diluted in synaptosomal RNA buffer (50 mM Hepes, pH 7.5, 125 mM NaCl, 100 mM sucrose, 2 mM K acetate, 10 mM EDTA) containing protease inhibitors (1 tablet/10 mL, Roche, Cat. N. 11836170001), 160 U/mL Superase-In (Invitrogen, Cat.N. AM2694), 160 U/mL Rnase-OUT (Invitrogen, Cat. N. 10777019), quickly pelleted at 20.000 g for 20 min and rinsed twice in 4 × volume of synaptosomal RNA buffer and spundown again 20.000 g for 20 minutes. They were then resuspended in 100 μL of synaptosomal RNA buffer and RNA was extracted with standard Trizol protocol (Chomczynski & Sacchi, 1987) using Qiazol lysis reagent (Qiagen Cat. N. 79306).

Validation of the Synaptosomes enrichment procedure was performed through Western Blot and qPCR. In the former case, the enrichment of NMDAR2B (abcam, cat. Num. ab65783) and the depletion of Histone H3 (abcam, cat. Num. ab1791), were evaluated and compared to α-Tubulin (Sigma-Aldritch, cat. Num. T9026) levels. In the latter case, complementary DNA (cDNA) was produced using the SuperScript IV Reverse Transcriptase (ThermoScientific, Cat. N. 18090010), following manufacturer’s instructions. cDNA was obtained from 100 ng of total brain extract RNA and total synaptosomal RNA and used for qPCR using SYBR Green PCR Master Mix (Qiagen, Cat. N. 208052) according to manufacturer’s instructions. A two-step program was run on a Biorad C1000 Touch cycler (Biorad) using 60° as annealing and extension temperature and generating a melting curve. The expression values of each synaptic and non-coding gene were normalized on H3F3B, coding for the histonic protein 3 B. (for details about the primers used see table below).

For the validation of ribosomal proteins increase with aging, the enrichments of RPL10A (abcam, cat. Num. ab226381), RPS6 (Cell signaling, cat. Num. #2217), RPL7 (Bethyl, cat. Num. A300-741A) and RPL15 (LS Bio, cat. Num. LS-C162700) were evaluated and compared to α-Tubulin (Sigma-Aldritch, cat. Num. T9026) levels, both in TH and SYN. The validation of the depletion of RNAs coding for ribosomal proteins from SYN by qPCR was done as described above. The expression values of each ribosomal gene were normalized on Ldhb, a transcript whose expression is stable in the two fractions analyzed across all time points (for details about the primers used see table below).

The validation of the differentially spliced regions by qPCR was done normalizing each junction with a reference exon as described in the text and didascalies (for details about the primers used see table below).

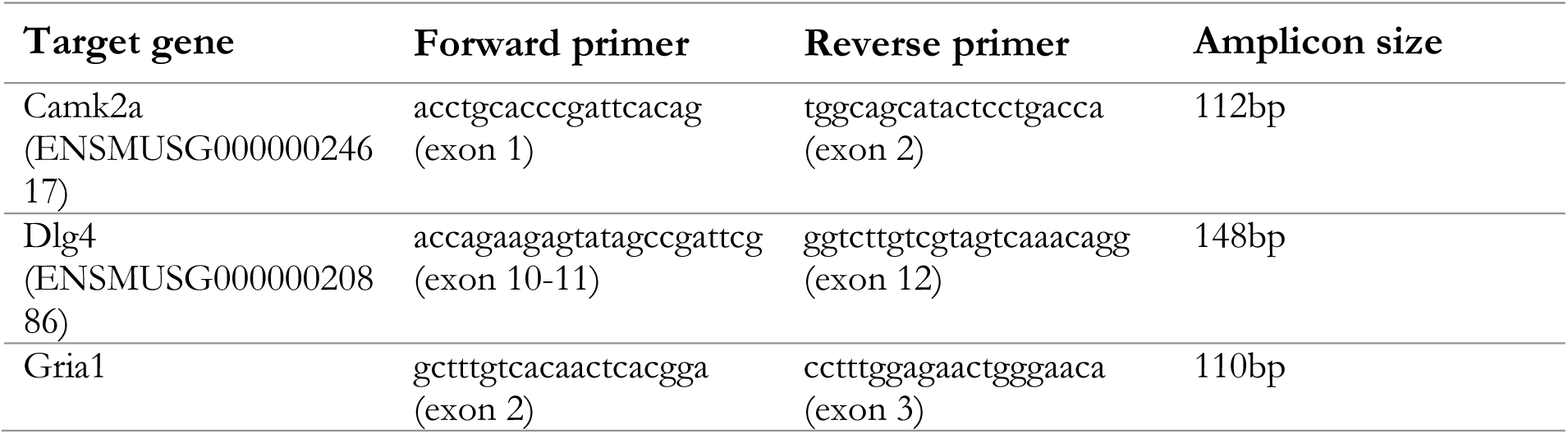
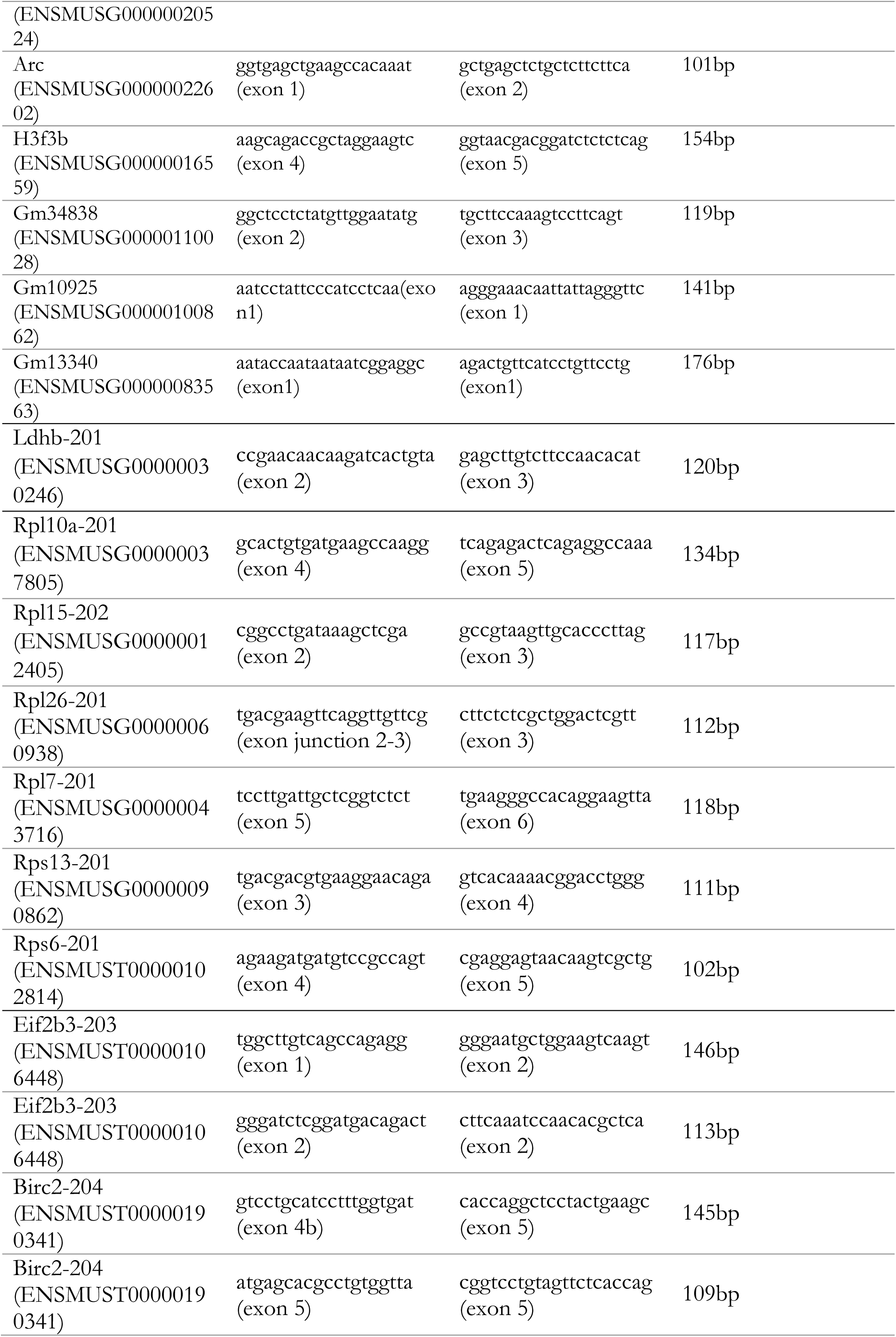
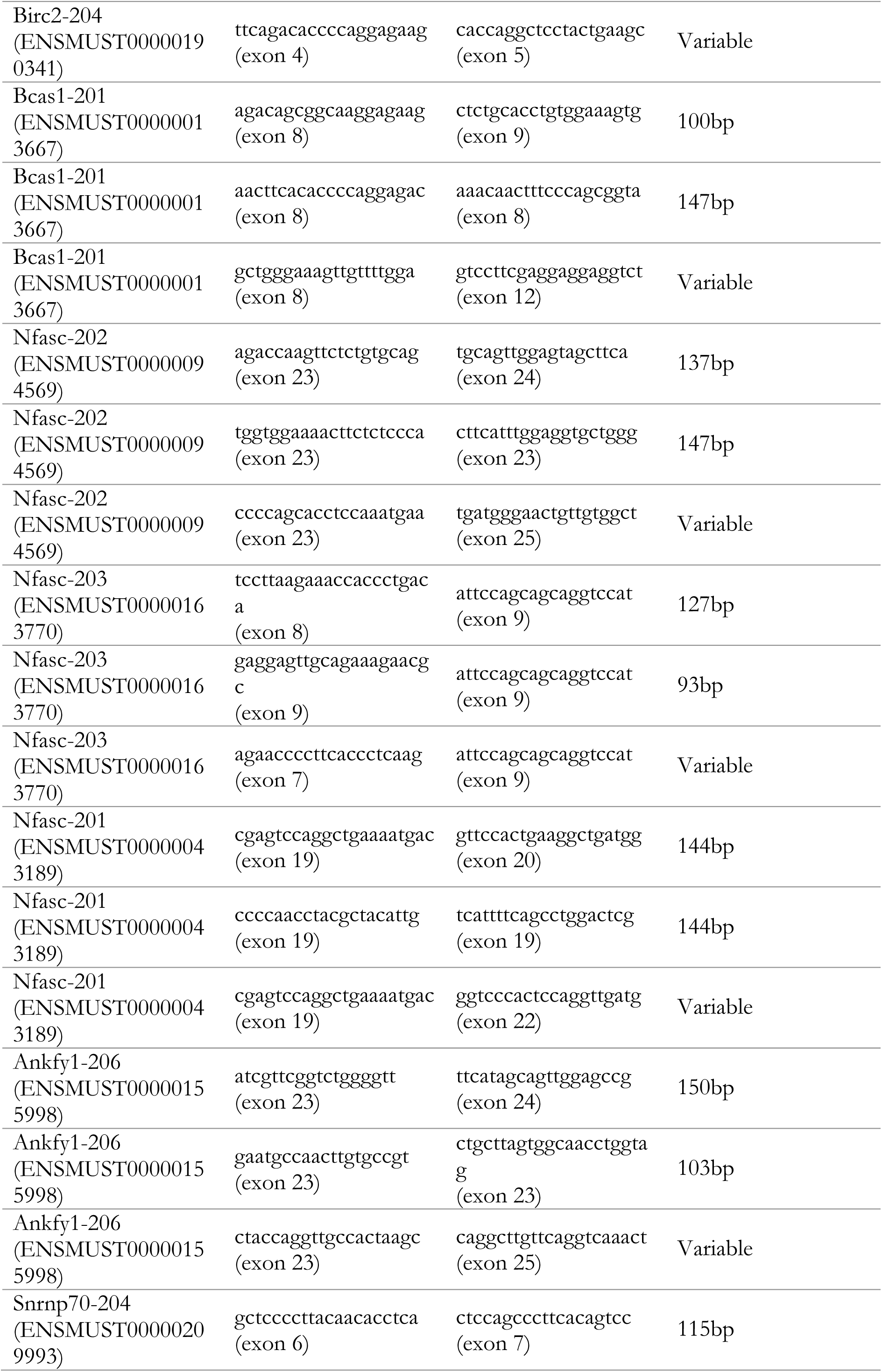
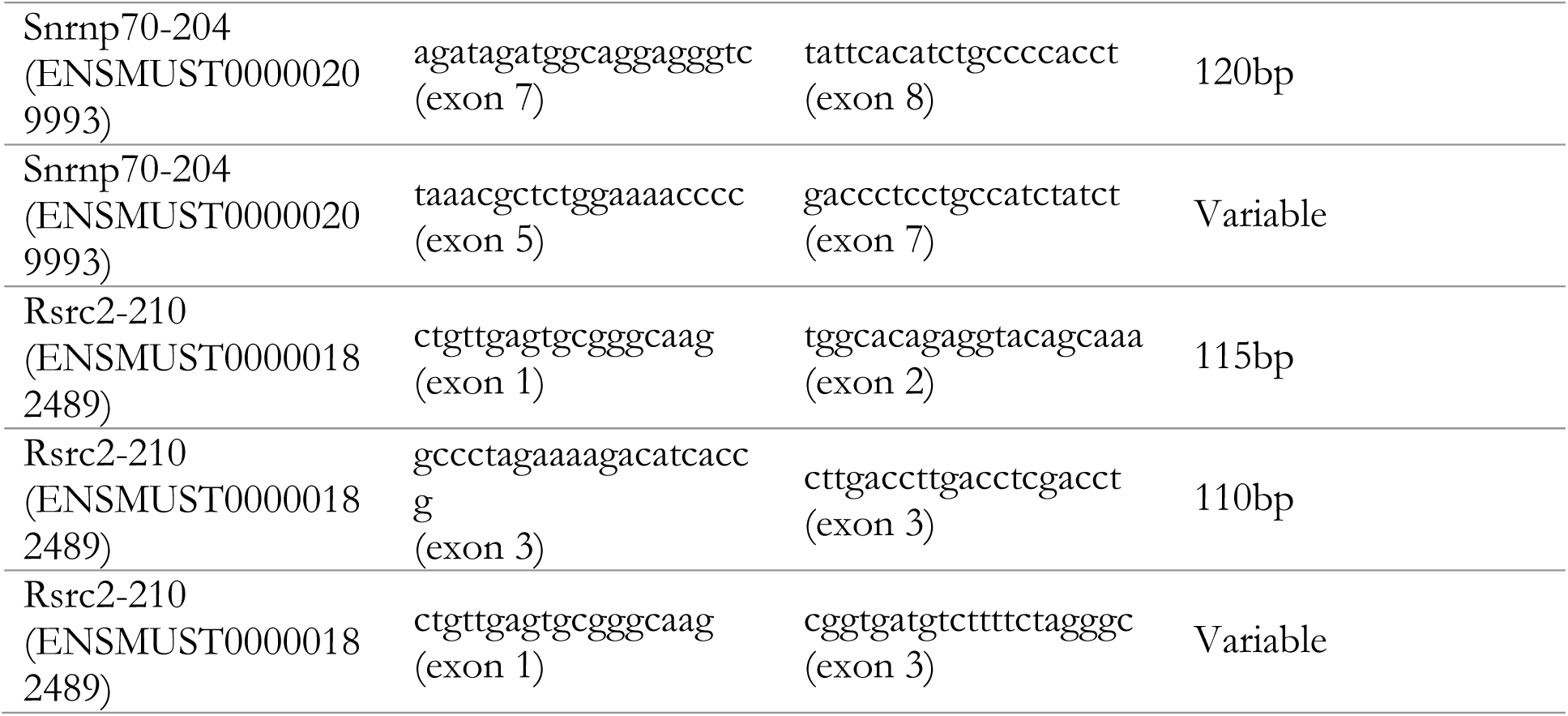

### Immunofluorescence of isolated synaptosomes

Isolated synaptosomes were plated on glass coverslips (10µg per coverslip) by centrifugation at 4000 rpm for 40 min at 4°C. Fixation was performed with 4% PFA in PBS (137 mM NaCl, 2.7 mM KCl, 10 mM Na2HPO4, 2 mM KH2PO4) at pH 7.5 for 10 min on ice followed by 35 min at room temperature. Coverslips were then washed briefly with PBS 2–3 times, and PFA was quenched with 100 mM NH4Cl for 25 min. They were then permeabilized and blocked in staining solution (PBS + 4% donkey serum, 2% BSA and 0.1% Triton X-100) for 30 min. Primary and secondary antibodies were applied in staining solution for 2 h and 1 h, respectively; washing between primary and secondary antibody incubation was 3 × 5 min, with PBS. After the secondary antibody incubation, cells were sequentially washed 3 × 5 min with PBS + 3 x 5 min high-salt PBS (PBS + 350 mM NaCl), and 3 x 5 min PBS to increase the stringency of the antibody staining. Finally, synaptosomes were embedded in ProLong glass antifade medium (Invitrogen, cat. Num. P36982) for imaging. The primary antibodies used were the following: Synaptophysin (Synaptic Systems, cat. Num 101 004), RPL7 (Bethyl, cat. Num. A300-741A) and rRNA (Y10b) (Novus biologicals, cat. Num. NB100-662). Coverslips were imaged with an AiryScan microscope (Zeiss). For unbiased, systematic processing of the images, an ImageJ macro was written to perform the analysis blind to age by keeping all parameters constant throughout. The synaptophysin channel (presynaptic vesicle pool marker) was used to identify the regions to analyze, and a constant threshold was applied and maintained for all images (to avoid bias across conditions). Individual synapses were selected and measured using the ImageJ particle analysis tool. Regions of interest (ROIs) were defined, added to the ROI manager, and automatically measured for the other two acquired channels (rRNA and RPL7). Fluorescence intensities and sizes were automatically saved to CSV files and further processed in R and GraphPad for statistical treatment.

### Isolation of ribosome associated RNAs from TH and SYN

Ribosome associated RNAs were purified using a sucrose cushion as previously described (Fusco et al., 2021). Briefly, pelleted synaptosomes or 50µL of TH were lysed with 1mL ribosome lysis buffer (20 mM Tris pH 7.4, 150 mM NaCl, 5 mM MgCl_2_, 24 U/mL TurboDNase, 100 µg/mL cycloheximide, 1% Triton-X-100, 1 mM DTT, RNasin(R) Plus RNase inhibitor 200 U/mL and 1x cOmplete EDTA-free protease inhibitor). TH samples were then homogenized with a syringe and a 0.4×20mm needle on ice and centrifuged for 10min at 10.000g at 4°C. TH and SYN were then loaded on 3mL sucrose solution (34% sucrose, 20 mM Tris pH 7.4, 150 mM NaCl, 5 mM MgCl_2_, 1 mM DTT, 100 µg/mL cycloheximide) in a thickwall ultraclear tube (Beckman, 344062) and centrifuged for 40 min at 4 °C at 55.000 rpm (367,600 × *g*) with a SW60 rotor. Ribosome-containing pellet was directly collected in Qiazol (Qiagen Cat. N. 79306) and processed for RNA extraction.

### RNA sequencing

RNA Sequencing was performed on the TH and SYN samples. Prior to that RNA integrity (RIN Score) and RNA concentration were estimated using the Bioanalyzer 2100 (Agilent) and the RNA 6000 Pico Kit and RNA 6000 Nano Kit for SYN and TH samples, respectively. Sequencing of RNA samples was performed using Illumina’s next-generation sequencing methodology (Bentley et al., 2008). In detail, total RNA was quantified and quality checked using Agilent 2100 Bioanalyzer in combination with an RNA 6000 Pico assay (both Agilent Technologies). Libraries were prepared from 10 ng of input material (total RNA) using the SMARTer Stranded Total RNA-Seq Kit - Pico Input Mammalian (Takara, Cat. N. 635006) following the manufacturer’s instructions. Quantification and quality checked of libraries was done using an Agilent 2100 Bioanalyzer instrument and a DNA 7500 kit (Agilent Technologies). Libraries were pooled and sequenced using a HiSeq 2500. System run in Rapid v2/51 cycle/single-end mode. Sequence information was converted to FASTQ format using bcl2FastQ v2.19.0.316. Per sample, the reads were mapped to the Mus musculus genome GRCm38 and respective annotation Release 85 [PMID: 29155950] using tophat2 v2.1 [PMID: 23618408] (parameters: --no-coverage-search --no-convert-bam --no-novel-juncs --no-novel-indels -T). Reads per gene were counted using featureCounts v1.5 [PMID: 24227677] (parameters: -s 0). Read counts were introduced into the statistical environment R in order to calculate RPMs (reads per million mappable reads) and RPKMs (reads per kilobase and million mappable reads). For calculation of RPKMs lengths of transcripts were taken from featureCounts output.

For the RNAs obtained by sucrose cushion, total RNA was quantified and quality checked using Tapestation 4200 instrument in combination with RNA ScreenTape (both Agilent Technologies). Libraries were prepared from 30-150 ng of input material (total RNA) using NEBNext Ultra II Directional RNA Library Preparation Kit in combination with NEBNext rRNA Depletion Kit v2 (Human/Mouse/Rat) and NEBNext Multiplex Oligos for Illumina (Unique Dual Index UMI Adaptors RNA) following the manufacturer’s instructions (New England Biolabs). Quantification and quality checked of libraries was done using an Agilent 4200 Tapestation instrument and a D1000 ScreenTape (Agilent Technologies). Libraries were pooled and sequenced in two NovaSeq6000 S1 100 cycle runs. System run in 101 cycle/single-end/standard loading workflow mode. Sequence information was converted to FASTQ format using bcl2fastq v2.20.0.422. Sequencing finished with an average of 39.6 million reads per sample. Mapping was done as described above.

For the alternative splicing analysis, a paired-end 2×150bp sequencing was performed using the Illumina HiSeq2500 System and the HiSeq Rapid SBS Kit v2. Sequence information was extracted in FASTQ format using Illumina’s bcl2FastQ 2.20.0.422, and processed raw reads were finally mapped to the Genome Reference Consortium Mouse Build 38 patch release 6 (GRCm38.p6) using Segemehl 0.3.4 in split-read mode (Hoffmann et al., 2009), obtaining SAM files. Using SAMtools 1.7 (H. Li et al., 2009b) sorted BAM files and BAI files were generated. Raw counts were obtained using the featureCounts function 1.6.0 of the Subread package (Liao et al., 2014) (- p and --minOverlap 10 –s 1 -t exon -g gene_id).

### Sample preparation for Mass spectrometry (MS)

Samples were brought to the same sample volume (125 μL) with MilliQ water, then a 2x lysis buffer was added in equal volume, to yield the samples in 250 μL volume containing 1% SDS, 100 mM HEPES pH 8.5 and 50 mM DTT. 4x acetone volume were added for protein precipitation. A maximum g of 3220 could be used during all steps to obtain protein pellets.

Samples were sonicated with 10 cycles (60 sec ON, 30 sec OFF) using the high energy mode in a Bioruptor (Diagenode) at 20 °C. The samples were then boiled (95 °C, 10 minutes) before being subjected to another 10 cycles in the Bioruptor. Iodoacetamide (IAA) was added to a final concentration of 15 mM and the samples incubated in the dark for 30 min at room temperature. Proteins were precipitated with 4 volumes equivalent of ice-cold acetone overnight at −20 °C. The following day, samples were centrifuged at 4 °C, 14.000 rpm for 30 min (Eppendorf 5804R benchtop centrifuge), then the acetone supernatant carefully removed. Protein pellets were washed twice with ice cold 80% acetone/20% water (500 μL each time), and centrifuged 4 °C, 14.000 rpm 10 min each wash. After the last wash, the acetone/water was removed and the pellets allowed to air dry at room temperature.

Protein pellets were then resuspended in digestion buffer (3 M Urea in 100mM HEPES, pH 8) at a concentration of 1 μg/μL and sonicated using a Bioruptor for 3 cycles (60 sec ON/30 sec OFF). Proteins were digested with Lys-C protease (Wako, Cat. N. 125-05061) 1:100 enzyme : protein ratio (4 hours, 37 °C, with shaking 1.000 rpm) then diluted with milliQ water (for a final concentration of 1.5 M Urea) and further digested with Trypsin (Promega, Cat. N. V5280) 1:100 enzyme : protein ratio (overnight, 37 °C, 650 rpm). Samples were acidified the following day by addition of 10% Trifluoracetic Acid (TFA) and then desalted with Waters Oasis® HLB μElution Plate 30μm (Cat. N. 186001828BA) in the presence of a slow vacuum. In this process, the columns were conditioned with 3×100 μL solvent B (80% acetonitrile; 0.05% formic acid) and equilibrated with 3x 100 μL solvent A (0.05% formic acid in milliQ water). The samples were loaded, washed 3 times with 100 μL solvent A, and then eluted into PCR tubes with 50 μL solvent B. The eluates were dried down with the speed vacuum centrifuge and dissolved at a concentration of 1 μg/μL in5% acetonitrile, 95% milliQ water, with 0.1% formic acid prior to analysis by LC-MS/MS.

### LC-MS/MS for Data Independent Analysis (DIA)

Peptides were separated using the nanoAcquity UPLC MClass system (Waters) fitted with a trapping (nanoAcquity Symmetry C18, 5μm, 180 μm x 20 mm) and an analytical column (nanoAcquity BEH C18, 1.7μm, 75μm x 250mm). The outlet of the analytical column was coupled directly to Q-Exactive HFX (Thermo Fisher Scientific) using the Proxeon nanospray source. Solvent A was water, 0.1 % formic acid and solvent B was acetonitrile, 0.1 % formic acid. The samples (approx. 1 μg) were loaded with a constant flow of solvent A at 5 μL/min onto the trapping column. Trapping time was 6 min. Peptides were eluted via the analytical column with a constant flow of 0.3 μL/min. During the elution step, the percentage of solvent B increased in a non-linear fashion from 0 % to 40 % in 60 min. Total runtime was 75 min, including clean-up and column re-equilibration. The peptides were introduced into the mass spectrometer via a Pico-Tip Emitter 360 μm OD x 20 μm ID; 10 μm tip (New Objective) and a spray voltage of 2.2 kV was applied. The capillary temperature was set at 300 °C. The RF ion funnel was set to 40%. Data from the TH samples were first acquired in DDA mode to contribute to a sample specific spectral library. The conditions were as follows: full scan MS spectra with mass range 350-1650 m/z were acquired in profile mode in the Orbitrap with resolution of 60000. The filling time was set at maximum of 20 ms with limitation of 1 x 106 ions. The “Top N” method was employed to take the 15 most intense precursor ions (with an intensity threshold of 4 x 104) from the full scan MS for fragmentation (using HCD normalized collision energy, 31%) and quadrupole isolation (1.6 Da window) and measurement in the Orbitrap (resolution 15000, fixed first mass 120 m/z).The peptide match ‘preferred’ option was selected and the fragmentation was performed after accumulation of 2 x 105 ions or after filling time of 25 ms for each precursor ion (whichever occurred first). MS/MS data were acquired in profile mode. Only multiply charged (2+ - 5+) precursor ions were selected for MS/MS. Dynamic exclusion was employed with maximum retention period of 30 s and relative mass window of 10 ppm. Isotopes were excluded. In order to improve the mass accuracy, internal lock mass correction using a background ion (m/z 445.12003) was applied. For data acquisition and processing of the raw data Xcalibur 4.0 (Thermo Scientific) and Tune version 2.9 were employed. For the DIA data acquisition the same gradient conditions were applied to the LC as for the DDA and the MS conditions were varied as described: Full scan MS spectra with mass range 350-1650 m/z were acquired in profile mode in the Orbitrap with resolution of 120.000. The default charge state was set to 3+. The filling time was set at maximum of 60 ms with limitation of 3 x 106 ions. DIA scans were acquired with 34 mass window segments of differing widths across the MS1 mass range. HCD fragmentation (stepped normalized collision energy; 25.5, 27, 30%) was applied and MS/MS spectra were acquired with a resolution of 30.000 with a fixed first mass of 200 m/z after accumulation of 3 x 106 ions or after filling time of 47 ms (whichever occurred first). Data were acquired in profile mode. All samples (both TH and SYN) had data acquired in DIA mode.

### Data analysis for DIA data

For library creation, the DDA data was searched using Pulsar (v. 1.0.15764.0.27599) (Biognosys AG, Switzerland). The data were searched against a species specific (Mus musculus) Swissprot database with a list of common contaminants appended, as well as the HRM peptide sequences. The data were searched with the following modifications: Carbamidomethyl (C) (Fixed) and Oxidation (M)/ Acetyl (Protein N-term) (Variable). A maximum of 1 missed cleavage was allowed and the identifications were filtered to satisfy FDR of 1 % on peptide and protein level.

Also for library creation, the DIA data was searched using Pulsar (v.1.0.15764.0.27599). The data were searched against a species specific (Mus musculus) Uniprot database with a list of common contaminants appended, as well as the HRM peptide sequences. The data were searched with the following modifications: Carbamidomethyl (C) (Fixed) and Oxidation (M)/ Acetyl (Protein N-term) (Variable). DpD (DDA plus DIA) libraries were then created by combining the spectral library created from the output of the DDA runs with the searched libraries using Spectronaut (v. 11, Biognosys AG, Switzerland)(Bruderer et al., 2015). This library contained 78.629 precursors, corresponding to 4.216 protein groups using Spectronaut protein inference. DIA data were then uploaded and searched against this spectral library. Relative quantification was performed in Spectronaut for each pairwise comparison using the replicate samples from each condition. The data (candidate table) and data reports were then exported to excel and further data analyses and visualization were performed with R-studio (version 0.99.902) using in-house pipelines and scripts.

### Data analysis

All the analyses were performed using R-Studio 1.1.456 (Leemans et al., 2009). Starting from the raw counts, normalized counts and log2 fold changes were estimated using DESeq2 1.22.2 (Love et al., 2014). To combine transcriptomic and proteomic data, p-values for differential expression at the protein and transcript level are combined in a meta-analysis (Fisher’s method) (Lury & Fisher, 1972). Fisher’s method combines extreme value probabilities from each test, commonly known as “p-values”, into one test statistic (X2) using the formula:

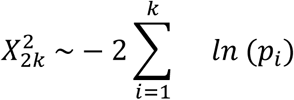

where pi is the p-value for the ith hypothesis test.

The visualization of sample’s similarity through Gene Enrichment has been evaluated by performing over representation analysis (ORA) using WebGestalt website (http://webgestalt.org/) (Liao et al., 2019). This website uses affinity propagation to cluster similar gene sets; it identifies the most enriched GO categories in a given dataset and provides the FDR-corrected p-value of the enrichment. The categories of the GO database (Ashburner et al., 2000; Carbon et al., 2017) that contain at least five genes were considered, and the p-values have been corrected for multiple comparisons through the FDR method described in (Benjamini & Hochberg, 1995). Visualization of Gene Enrichment was achieved using ReviGO software (http://revigo.irb.hr/). ReviGO allows to obtain a flexible reduction in size for large user-supplied lists of overlapping GO categories, and visualize the remaining GO terms in a two-dimensional space which reflects the terms’ semantic interrelations.

Classification of genes as coding or non-coding was done using Ensembl BioMart (https://www.ensembl.org/biomart/) annotation version GRCm38.p6.

Principal Component Analysis and Heatmap representation of their hierarchical clustering has been performed using DESeq2’s internal functions and the packages ComplexHeatmap 2.0.0, pcaMethods 1.76.0 and ggpubr 0.2. Density plots have been drawn using the packages ggplot2 3.1.1 and ggpubr 0.2.

Generally Applicable Gene-set Enrichment for Pathway Analysis (GAGE) was performed using gage 2.34.0 and GO.db 3.8 packages (Luo et al., 2009b). GAGE is a published method for gene set (enrichment or GSEA) or pathway analysis. To test whether a gene set is significantly correlated with a phenotype or an experiment condition, the fold changes of gene expression level in the experiment condition (or phenotype) are compared to a control condition. Then, a two-sample t-test is applied to verify whether the mean fold changes of a target gene set is significantly different from that of the background set (the whole gene set of the microarray).

In ribosome profiling experiments, Translational Efficiency (TE) is generally defined as the ratio between the Ribosome Protected Fragments (RPFs) over mRNA counts. Since we were dealing with whole transcripts associated with ribosomes, we calculated SYN TE as the ratio between ribosome associated transcripts (SC) in SYN and TH. When examining a single compartment (either SYN or TH), we calculated TE as the ratio between ribosome associated transcripts (SC) and all transcripts in that compartment (input).

### Identification of Differentially Expressed Junctions (DEJs)

DEJs were identified using the software DIEGO (Doose et al., 2018) and LeafCutter (Y. I. Li et al., 2018). Gene models and the coverage at genomic intervals of interest was visualized using the software IGV 2.5.2 (Robinson et al., 2011; Thorvaldsdóttir et al., 2013) and the R packages rtracklayer 1.44.0 (Lawrence et al., 2009), GenomicRanges 1.36.0 and GenomicFeatures 1.36.1 (Lawrence et al., 2013) and ggbio 1.32.0 (Yin et al., 2012).

## ACKNOWLEDGMENTS

The Core Facilities and Services CF Next generation sequencing (Cornelia Luge and Ivonne Goerlich), CF Life science computing and CF proteomics of the FLI are gratefully acknowledged for their technological support. Dr Mario Baumgart is gratefully acknowledged for the help with internal animal permissions.

## DATA AVAILABILITY

The RNA-seq datasets here presented are available on GEO: GSE281456, GSE281455 and GSE146636. The proteomics dataset is available on MassIVE at the following link: https://massive.ucsd.edu/ProteoSAFe/private-dataset.jsp?task=b2cf624f9229489eb9f47b22b7e9244f. Password for reviewers is FLIReviewer_2024.

## CONFLICTS OF INTEREST

The authors declare no conflicts of interests.

## AUTHOR CONTRIBUTIONS

CC, MU, SH, AC project design CC,MU,WD,KJ,MG,KR,MG: data production CC,MU,EF,AC: data analysis

CC,MU,EF,SH,AC: data interpretation SH,AC: supervision and coordination

CC,MU,AC: drafting the manuscript

All authors contributed to the final version of the manuscript

## Notes

### Competing Interest Statement

The authors have declared no competing interest.

